# Lymphatic constraint of germinal centers optimizes protective antibody responses

**DOI:** 10.1101/2025.11.03.686378

**Authors:** Tenny Mudianto, Gabrielle R. Berman, Kimberly Zaldana, Luan Firmino Cruz, Marta Schips, Niklas Schwan, Ochapa Ibrahim, Vanessa Cristaldi, William S. Foster, Lina Fikri, Sabrina Solis, Han Chen, Michael Cammer, Ramin Sedaghat Herati, Sergei B. Koralov, Andrew David Miller, Michael Meyer-Hermann, Carla R. Nowosad, Amanda W. Lund

## Abstract

Immunization strategies are central to pathogen control, where efficacy relies on antigen uptake, distribution, persistence, and inflammatory context. We recently demonstrated that dermal lymphatic capillaries regulate antigen presentation in lymph nodes (LN) by restraining fluid and virion transport following vaccinia virus (VACV) infection by skin scarification. Concurrently, a perifollicular, LN lymphangiogenic response encapsulates expanding B cell follicles. Given the important role of antigen transport and uptake on humoral immunity, we tested the hypothesis that lymphatic remodeling in the skin and LNs regulates germinal center (GC)–dependent antibody responses during infection. Using a model of lymphatic-specific VEGFR2 inhibition, we found that inhibiting viral-induced lymphatic remodeling in skin and LNs prompted significant GC expansion but paradoxically decreases protective VACV-specific class-switched antibodies. While the larger GC responses appeared structurally normal, they failed to support a proliferative burst consistent with clonal selection. Mathematical modeling revealed that this disconnect between GC size and function arises from impaired productive T follicular cell interactions in larger GC volumes and consistent with this finding, the optimal GC size was evolutionarily conserved across diverse mammals. Finally, we found that the presence of virus in the LN initiates these changes in GC function, inhibits LN lymphangiogenesis, increases B cell follicle size, and reduces selection efficiency. Therefore, protecting LN lymphatic vessels from virus-induced interferons rescues perifollicular lymphatic growth and follicle size, indicating that LN lymphangiogenesis directly constrains the follicular response. In summary, this study underscores the central role of lymphatic remodeling in compartmentalizing antigen and inflammatory signals to optimize GC fitness and protective antibody responses.

## INTRODUCTION

Conventional immunization strategies remain a mainstay of efforts to prevent disease transmission and infection-associated pathology, and yet are often insufficient to generate broadly neutralizing antibodies that protect against pathogenic rechallenge. While significant vaccine design efforts are focused on the nature of the antigen and adjuvant, vaccine biodistribution – the transport and persistence of vaccine components in the host - is also likely a key determinant of vaccine efficacy. Consistently, the orthopoxvirus vaccinia virus (VACV) generates durable humoral and cellular immunity^2^ superior to other routes of dissemination (e.g. intradermal, intramuscular), when delivered epicutaneously to superficially injured skin (skin scarification)^1^. Slow, sustained immunization in rhesus monkeys resulted in more robust T follicular helper (T_FH_) cell responses and higher titers of neutralizing, germinal center (GC)-dependent antibodies than conventional bolus immunization^3^. Furthermore, the organ distribution, expression kinetics, and therapeutic outcomes of lipid nanoparticles, widely used to deliver mRNAs, depend upon their size and route of administration^4^. Therefore, understanding the endogenous mechanisms that regulate pathogen and antigen transport may provide parameters to further optimize protective antibody generation from new and existing vaccines.

Peripheral tissue lymphatic capillaries are blunt-ended structures that facilitate the transport of interstitial fluid, molecular species, and leukocytes to secondary lymphoid organs^5^ and regulate the distribution of cell-free and cell-bound antigen. Naïve lymphatic capillaries maintain specialized, discontinuous junctions, termed buttons, that permit passive paracellular transport^6^, and lymph node (LN) lymphatic endothelial cells (LECs) provide survival cues for antigen scavenging subcapsular sinus (SCS) macrophages, shape chemokine gradients, and filter molecular species that enter the conduit system^7^. Rather than being static structures, however, we recently demonstrated that dermal lymphatic capillaries actively tune their transport function to restrict the passive flux of fluid and infectious virions from the skin to LNs^8,9^. The constraints placed on passive transport by the peripheral lymphatic vasculature optimize dendritic cell (DC) migration and CD8^+^ T cell priming, indicating a functional role for active, infection-induced lymphatic remodeling in shaping anti-viral immune responses. In addition to cellular immunity, anti-viral antibody responses depend on lymphatic transport to LNs^9^.

The germinal center (GC) is a spatially regulated immune niche that sits just under the SCS and is specialized for the interaction of B cells with antigens and helper T cells. Antigen transported by afferent lymphatic vessels is scavenged by SCS macrophages and transferred to follicular dendritic cells (FDC) to initiate a GC, where rounds of mutation and selection expands B cells with improved binding of their surface B cell receptor (BCR) to antigens, and in turn, improved affinity of their secreted antibody^10^. B cell activation and assignment to the GC fate also supports the process of isotype-switching to generate specialized isotypes such as IgG, the isotype favored for systemic neutralization of viruses. Rather than imposing strict affinity thresholds, GCs are relative structures^11^ that create the framework for competition to expand the best performing B cells in the pool at any given time through T cell-dependent clonal expansion ^10,12^. Iterative rounds of mutation, selection, and expansion causes successful clones with improved affinity to predominate the GC, secreted antibody, and memory B cell response over time^10,13,14^. In addition to the changes in lymphatic transport that would alter the delivery of GC activating signals, VACV infection expands lymphatic structures into the interfollicular areas of the LN parenchyma that appear to encapsulate activating B cell follicles^15,16^, though the functional consequence of this growth remains unknown. Since small changes to the initiation, maintenance, and selection dynamics within each individual GC can alter GC competitiveness and have large impacts on downstream antibody-dependent responses, here we seek to investigate the functional significance of lymphatic remodeling on GC-dependent antibody responses.

In this study, we test the hypothesis that viral-induced lymphatic remodeling, including both peripheral capillary zippering and LN lymphangiogenesis, shape GC dynamics and thereby regulate protective antibody output. To test this hypothesis, we employ a genetically engineered conditional mouse model in which vascular endothelial growth factor receptor (VEGFR) 2 is specifically deleted in lymphatic vessels, inhibiting VACV-induced changes in lymphatic transport^8^ and the burst of perifollicular LN lymphangiogenesis that precedes GC onset. We demonstrate that the loss of perifollicular lymphatic structures in the LN associates with expansion of the B cell follicle and GC reaction, but diminished antibody abundance and reduced protection against VACV rechallenge. The surprising negative association between GC size and antibody output depended on a failure to select and expand clones within individual GCs, explained by a size-dependent reduction in the probability of amassing positive selection signals via T_FH_ interactions. Importantly, the lymphatic vasculature regulated GC size and subsequent selection in two ways, first by restraining viral transport from the periphery^8^, which was sufficient to replicate the VEGFR2-dependent phenotype, and second by maintaining a local inflammatory environment that permitted rapid lymphatic growth. Indeed, loss of lymphatic-intrinsic type I interferon (IFN) signaling was sufficient to rescue perifollicular lymphangiogenesis and subsequently constrain B cell follicle size even in the presence of viable virus. We therefore present a novel mechanism through which lymphatic remodeling sets a fundamental size constraint on the GC response that is conserved across mammals and is required for optimal antibody production and protective immunity.

## RESULTS

### A perifollicular lymphangiogenic response precedes germinal center formation following viral infection

Vaccinia (VACV) infection administered by skin scarification induces VEGFR2-dependent peripheral lymphatic zippering that reduces fluid flow and passive viral dissemination but optimizes dendritic cell (DC) trafficking and early CD8 T cell priming^8^. These observations indicated that active mechanisms of lymphatic remodeling in response to viral infection play a key role in optimizing the anti-viral cellular response but whether these changes in transport also impact on humoral responses, which similarly depend on lymphatic transport^9^, remains unknown. We therefore began to investigate the dynamics of the humoral response within the draining superficial cervical LN following VACV infection by skin scarification. Upon infection by scarification, VACV remains local at the site of infection with peak titers from days 3-5 post infection, does not infect the draining LN, and is cleared by day 15^9,17^. Stromal components of the LN are known to undergo remodeling as a function of peripheral tissue inflammation^18^, including LN lymphatic vessels. Indeed, in draining LNs we observed lymphatic growth that appeared to invade into the LN cortex encapsulating the activating B cell follicle as early as 5 days post-infection and continued to surround follicles containing active GCs 15 days post-infection (**Figure 1A**). By pulsing with bromodeoxyuridine (BrdU) at five-day intervals post infection, we observed a peak in LEC proliferation from days 0-5 post-infection (**Figure S1A and 1B**) with increased LEC numbers continuing to accumulate over time in concert with LN expansion (**Figure S1B**). This burst in lymphatic proliferation preceded the expansion of other LN stromal subsets including fibroblastic reticular cells (FRC) and blood endothelial cells (BEC) (**Figure S1C and D**) as well as the activation of germinal center (GC) B cells (**Figure 1A and C**). To investigate the nature of this lymphangiogenic response at higher resolution, we performed single cell RNAseq analysis of sorted LN endothelial cells (CD45^-^CD31^+^) from days 0, 7, 15, and 25 post-infection. We subset on LECs (*Prox1*^+^*Lyve1*^+^) (**Figure S1E and F**) and reclustered the data to define LEC subsets in resting and inflamed LNs. Data from each time point was integrated with Harmony^76^ before visualization using uniform manifold approximation projection (UMAP) (**Figure 1D**). Unbiased clustering of the data identified five LEC clusters aligned with the literature^19^, including ceiling (*Ackr4*^+^*Foxc2*^+^*Lyve1*^-^), floor (*Lyve1*^lo^*Cd274*^+^*Madcam1*^+^*Ccl20*^+^), Marco+ (*Lyve1*^+^*Marco*^+^*Cd274*^+^) and PTX3+ (*Lyve1*^+^*Ptx3*^+^*Cd274*^-^) homeostatic subsets and an inflamed population expressing markers consistent with an inflamed state induced by oxazalone treatment^19^ (e.g. *Ccl2*, *Lgmn*, *Samhd1, Cxcl9*, *Ifit1*) (**Figure S1G and H**). These inflamed LECs also exhibited expression of Ptx3 and Marco indicating that likely arose from these homeostatic subsets (**Figure S1G**). When assessed over time, we found that there was significant enrichment for PTX3^+^ and inflamed LECs relative to other subsets at effector time points, 7-15 days post-infection (**Figure 1E and F**). We sought to validate these changes by flow cytometry, where PD-L1 and LYVE1 can be used to distinguish four LEC states consistent with the transcriptional clusters observed in single cell, including LYVE1^+^PD-L1^-^ (PTX3-like), LYVE1^+^PD-L1^+^ (Marco-like), LYVE1^-^PDL1^+^ (Floor), and LYVE1^-^PDL1^-^ (Ceiling) (**Figure 1G and S1G**). Here we observed enrichment of both the PTX3 (LYVE1^+^PD- L1^-^) and Marco (LYVE1^+^PD-L1^+^) LECs relative to the floor and ceiling subsets at day 5 post infection (**Figure 1H and I**), indicating that these subsets likely contribute to the first proliferative lymphangiogenic response.

**Figure 1.**
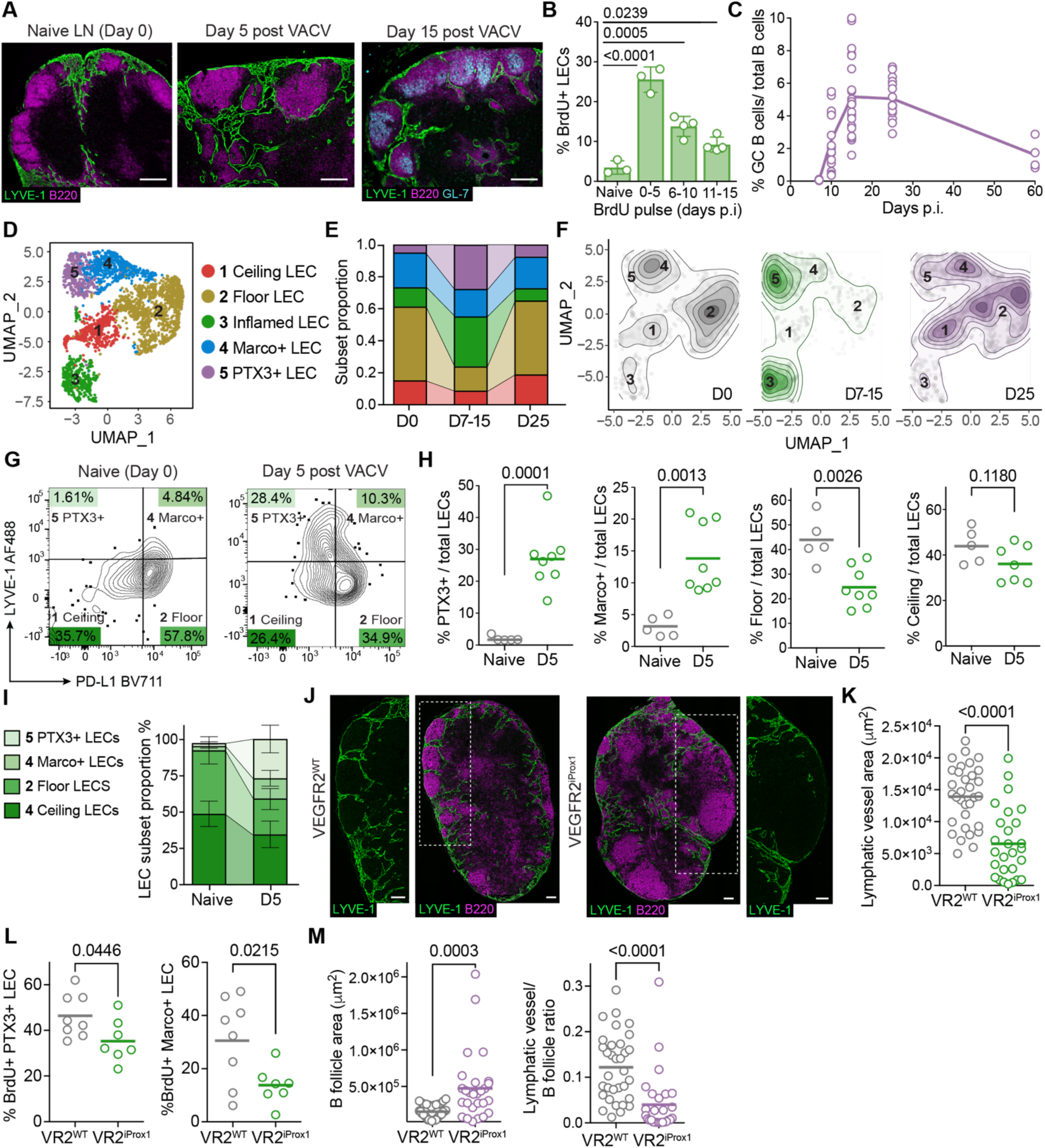
A perifollicular lymphangiogenic response associates with B cell follicle expansion. (**A**) Representative immunofluorescence images of lymphatic vessel and B cell follicle organization in draining lymph nodes (dLNs) from C57Bl/6 wildtype mice prior to, or at indicated time points post vaccinia virus (VACV) infection (by scarification). Green, LYVE-1; magenta, B220; cyan, GL-7. Scale bars = 200 μm. *p* values calculated using one way ANOVA. (**B**) Frequency of BrdU⁺ LN lymphatic endothelial cells (LECs; CD45⁻CD31⁺gp38⁺) following BrdU pulse-labeling at indicated five-day windows post VACV infection. Mice analyzed 5 days after BrdU labeling. Each point is one mouse; *p* values calculated using two-tailed Student’s *t* test. (**C**) Frequency of GC B cells (CD19^+^B220^+^GL-7⁺CD38^-^) of total B cells (B220^+^) over time post VACV scarification. Each point is one mouse; line connects mean values of each time point. (**D**) UMAP projection of scRNA-seq transcriptomic profiles of LN LECs (*Prox1*^+^) sorted from C57BL/6 WT mice collected at multiple time points post VACV infection and colored by Seurat cluster. Superficial cervical LNs were pooled for the analysis; naïve = 10 mice, days 7, 15, 25 = 5 mice each. (**E**) Proportional distribution of Seurat clusters over time, colored as in (D). Day 7 and 15 post infection time points combined. (**F**) Contour plot of UMAP in (D) split by time point. (**G**) Representative flow cytometry plots of LN LECs (CD45⁻CD31⁺gp38⁺) from naïve and dLNs at day 5 post VACV infection. Quadrants define four LEC subsets: [1] LYVE1^hi^ PD-L1⁻; [2] LYVE1^hi^ PD-L1⁺; [3] LYVE1^lo^ PD-L1⁺; [4] LYVE1^lo^ PD-L1^-^. (**H**) Quantification of LEC subsets defined in (G) as a proportion of total LECs in naïve versus day 5 dLNs. Each point is one mouse; data are from two independent experiments, *p* values calculated using two-tailed Student’s *t* test. (**I**) Proportional distribution of LEC subsets over time, from (H). (**J**) Representative immunofluorescence images of lymphatic vessels and B cell follicles in dLNs of VEGFR2^WT^ (VR2^WT^) and VEGFR2^iProx1^ (VR2^iProx1^) mice at day 7 post VACV infection. White box indicates high magnification inset. Green, LYVE-1; magenta, B220. Scale bars = 200 μm. (**K**) Quantification of perifollicular lymphatic vessel area surrounding B cell follicles in dLNs of VEGFR2^WT^ and VEGFR2^iProx1^ mice at day 6–7 post VACV infection. Each point is one follicle. Data are from two independent experiments (n=8 mcie, VEGFR2^WT^; n=6 mice, VEGFR2^iProx1^). Lines indicate mean; *p* values were calculated using two-tailed Student’s *t* test. (**L**) Frequency of proliferating BrdU⁺ LECs in expanding LEC subsets (1 and 2) in VEGFR2^WT^ and VEGFR2^iProx1^ dLNs pulsed with BrdU from day 0–5 post VACV infection. Each point is one mouse; data are from two independent experiments (n=8 mice, VEGFR2^WT^; n=7 mice, VEGFR2^iProx1^). Lines indicate mean; *p* values were calculated using two-tailed Student’s *t* test. (**M**) Quantification of B follicle area and the ratio of lymphatic vessel area per follicle size in dLNs from (K). Each symbol represents one follicle. Data are from two independent experiments (n=8 mice, VEGFR2^WT^; n=6 mice, VEGFR2^iProx1^). Lines indicate mean; *p* values were calculated using two-tailed Student’s *t* test.

While previous reports describe a VEGFA/VEGFR2-dependent perifollicular lymphangiogenic response^16^, the functional significance of the interrelationship between lymphatic growth and B cell responses remains unknown. We therefore asked whether we could use lymphatic-specific VEGFR2 knockout mice to investigate the impact of perifollicular lymphangiogenesis on GC induction and function post VACV infection. We therefore used the Prox1:Cre^ERT2^ to drive the inducible deletion of *Vegfr2*^fl/fl^ (VEGFR2^iProx1^) specifically in lymphatic vessels in the adult prior to infection. Consistent with the literature^16^ and our prior work^8^, we found that lymphatic-specific deletion of VEGFR2 (**Figure S1I and J**) was sufficient to blunt the infection-induced lymphangiogenic response (**Figure S1K and L**) relative to controls (VEGFR2^WT^). This decrease in proliferation associated with a loss the perifollicular lymphangiogenic response (**Figure 1J and K**), which associated with the reduction of LYVE1^+^PDL1^-^ and LYVE1^+^PDL1^+^, which include the PTX3^+^ and MARCO^+^ LEC subsets, respectively (**Figure 1L**). Surprisingly, this reduction in LN lymphangiogenesis led to a significant expansion of B cell follicle area in the first week of infection (**Figure 1M**), with B cell follicles lacking an encapsulating lymphatic network exhibiting the largest cross-sectional area (**Figure J and M**). These data pointed toward a functional interplay between lymphatic remodeling and the early B cell response to infection.

### Lymphatic remodeling constrains the germinal center response and enhances the production of protective antibodies

Based on the functional association between LN lymphangiogenesis and the structure of the early B cell response, we asked whether VEGFR2-dependent lymphatic remodeling impacted on the infection-induced GC response. As seen at early time points, we found a increase in the magnitude of the GC response 15 days post-infection (**Figure S2A and 2A-B**). Despite their size, however, GCs formed in this context appeared structurally similar, proportionately scaling their light and dark zones (LZ and DZ, **Figure 2C**) and maintaining an equal density of follicular helper T cells, (T_FH_, **Figure S2B and 2D**). Surprisingly, however, the enhanced size of the GC response did not translate to more productive antibody responses. When normalizing to total concentration of circulating immunoglobulin (Ig, **Figure 2E**), we found a reduction in total VACV-specific antibodies 15 days post-infection (**Figure 2F**) that was largely the result of the reduction in IgG2b/c isotypes, which are GC-dependent and critical for systemic protection against viral pathogens (**Figure 2G**). This deficiency in anti-VACV IgG2b/c antibody production persisted through memory time points (**Figure S2C-F**), indicating that despite their size, GCs induced in mice with impaired lymphatic remodeling were less functional.

**Figure 2.**
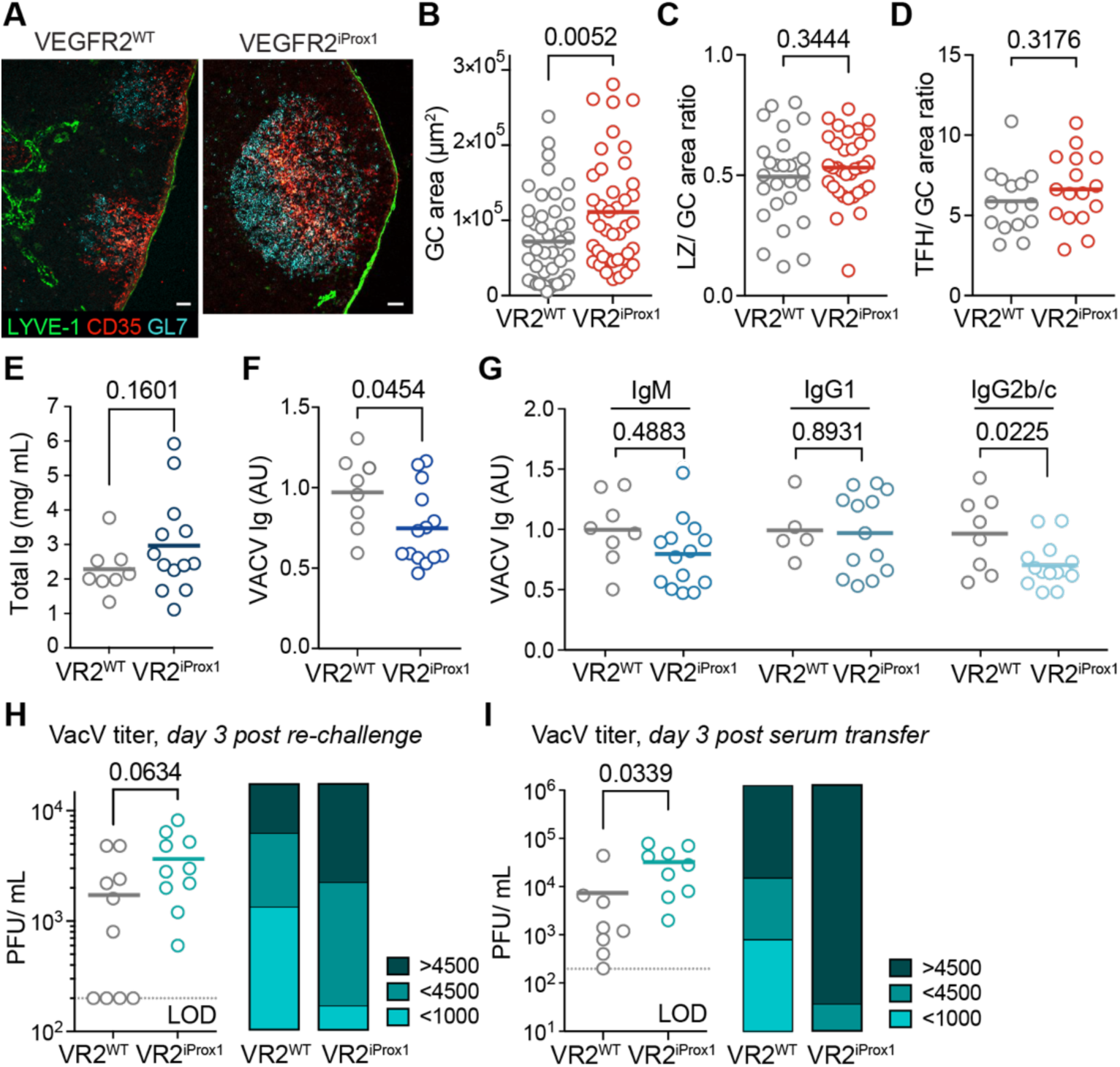
VEGFR2-dependent lymphatic remodeling enhances the formation of virus-specific antibodies. (**A**) Representative immunofluorescence images of germinal center (GC) zonal organization in draining lymph nodes (dLN) of VEGFR2^WT^ (VR2^WT^) and VEGFR2^iProx1^ (VR2^iProx1^) mice at day 15 post vaccinia virus (VACV) infection (by scarification). Green, LYVE-1; cyan, GL-7; red, CD35. Scale bars = 50 μm. (**B-D**) Quantification of GC area (B), ratio of light zone (LZ, CD35) area to total GC area (C), and ratio of follicular helper T cells (T_FH_, CD3e) area to GC area (D) in dLNs of VEGFR2^WT^ and VEGFR2^iProx1^ mice at day 15 post VACV infection. Each point is one GC. Data in (B and C) are from three independent experiments (n=8 mice, VEGFR2^WT^; n=6 mice, VEGFR2^iProx1^), and (D) is from one experiment (n=2 mice, VEGFR2^WT^; n=4 mice, VEGFR2^iProx1^). p values were calculated using two-tailed Student’s *t* test. (**E-G**) Serum antibody measurements in VEGFR2^WT^ and VEGFR2^iProx1^ mice at day 15 post VACV infection, assessed by ELISA. Total serum Ig concentration (E), normalized absorbance unit (AU) of serum VACV-specific IgG (F), and VACV-specific antibodies of different isotypes (IgM, IgG1, and IgG2b/c) (G). Data in (F and G) are normalized by the median values of each VEGFR2^WT^ control group. Each point is one mouse; lines indicate mean. Data are from three independent experiments (n=8 mice, VEGFR2^WT^; n=13 mice, VEGFR2^iProx1^); *p* values were calculated using two-tailed Student’s *t* test. (**H-I**) Viral titers (plaque-forming units, PFU) in infected ears of VACV-immune VEGFR2^WT^ and VEGFR2^iProx1^ mice collected at day 3 post VACV rechallenge (H) and in C57BL/6 WT mice treated with serum from VEGFR2^WT^ or VEGFR2^iProx1^ donors prior to rechallenge. Titers quantified at day 3 post VACV rechallenge (**I**). Dotted lines indicate the limit of detection (LOD). Stacked bar graphs (right) show proportion of mice with low, medium and high viral titers (>4,500 (high), <4,500 (medium), or <1,000 (low) PFU/mL). Each point is one mouse. Data are from two independent experiments; *p* values were calculated using two-tailed Student’s *t* test.

To test whether this reduction of VACV-specific antibodies had a functional significance, we performed a homologous rechallenge experiment. Memory VEGFR2^iProx1^ and VEGFR2^WT^ mice, 40 days post-VACV infection, were treated with both CD4 and CD8 depleting antibodies to remove protective memory T cells prior to re-infection with VACV on the uninfected, contralateral ear of the mouse (**Figure S2G and H**). Viral titers were evaluated three days post re-infection, where T cell-independent protection was observed in wildtype C57Bl/6 mice (**Figure S2I**). Three days post re-challenge, we found that VEGFR2^WT^ mice were more likely to be completely protected from re-infection, whereas all VEGFR2^iProx1^ mice had detectable virus in their skin (**Figure 2H**). To then evaluate the functional capacity of circulating antibodies more specifically, we performed passive serum transfer experiments. Convalescent serum from either VEGFR2^iProx1^ or VEGFR2^WT^ VACV mice was administered to naïve C57Bl/6 recipients one hour prior to VACV infection (**Figure S2J**). Three days post-infection, we again observed significantly higher viral titers in the skin of mice receiving VEGFR2^iProx1^ serum (**Figure 2I**), indicating reduced protective efficacy compared to serum from VEGFR2^WT^ donors. These data thus indicated that despite the expansion of the GC niche itself, the protective capacity of VEGFR2^iProx1^ GC-derived antibodies was inferior to that of wildtype controls, demonstrating a direct functional relationship between active lymphatic remodeling, GC size, and anti-viral humoral immunity.

### B cell selection is deficient in germinal centers lacking lymphatic constraint

These data clearly showed that despite the seemingly normal GC architecture, the efficiency of the GC response was compromised in the absence of VEGFR2-dependent lymphatic remodeling. Therefore, we sought to understand how the dynamics of the GC response might be altered in the context of the VEGFR2^iProx1^ mice. To do this, we performed scRNAseq analysis of sorted and then pooled GC B cells (B220^+^CD38^-^GL7^+^), plasma cells (PC; B220^lo^CD138^+^), and T_FH_ (TCRβ^+^CD4^+^PD1^+^CXCR5^+^) from three VEGFR2^WT^ and three VEGFR2^iProx1^ mice 25 days post-infection (**Figure S3A**). After sub-setting on B cells (*Cd19*) we performed unbiased clustering and UMAP visualization to identify four clusters^10,20,21^, consisting of plasma cells (PC; *Sdc1*, *Xbp1*), and GC B cells (GC; *Cd19*, *Bcl6*) (**Figure 3A and S3B**), which were equally distributed between both groups (**Figure 3B**). The GC cluster included transcriptionally distinct LZ subsets (*Cd86*, *H2-Oa*), DZ subsets (*Foxo1*, *Mki67*), and pre-memory B cells (Ccr6, Sell) (**Figure 3D and S3B**). LZ and DZ cluster annotations were further validated by cell cycle analysis, where DZ cells were cycling (S and G2/M phases) and LZ cells were in G1 arrest. Activated LZ cells (*Myc*^high^) exhibited transcriptional profiles indicative of recent BCR signaling (**Figure S3C and D**). Clustering the data at a higher resolution revealed eight clusters that highlighted different stages of the LZ to DZ cycle (**Figure 3C and S3E and F**). Cell cycle analysis mapped expected patterns of proliferation with LZ B cells in G1 transitioning through S phase to G2/M phase in the DZ with no significant differences between groups (**Figure S3C and D**). While a principal component analysis (PCA) of the individual samples showed no significant differences in the transcriptional profiles of the clusters between groups (**Figures S3G**), analysis of the proportion of each cluster represented per group showed a significant shift, where GC B cells extracted from VEGFR2^iProx1^ were significantly diminished in clusters 5 and 6, which represented the transitory *mki67*^+^ LZ/DZ and an early *mki67*^+^ proliferating DZ population, respectively (**Figure 3D-F**). These data began to point to a defect in clonal selection, rather than intrinsic differences in GC B cell transcriptional programming. Clonal selection occurs when GC B cells compete for signals from T_FH_ in the LZ to facilitate their full activation and transition to the DZ to undergo proliferation. Since the magnitude of B cell activation is known to drive DZ proliferation potential^10,22^, a specific reduction of highly activated recent LZ emigrants could result in a deficit in the signals necessary to drive strong proliferation and selection and impact the generation of high-affinity antibody responses.

**Figure 3.**
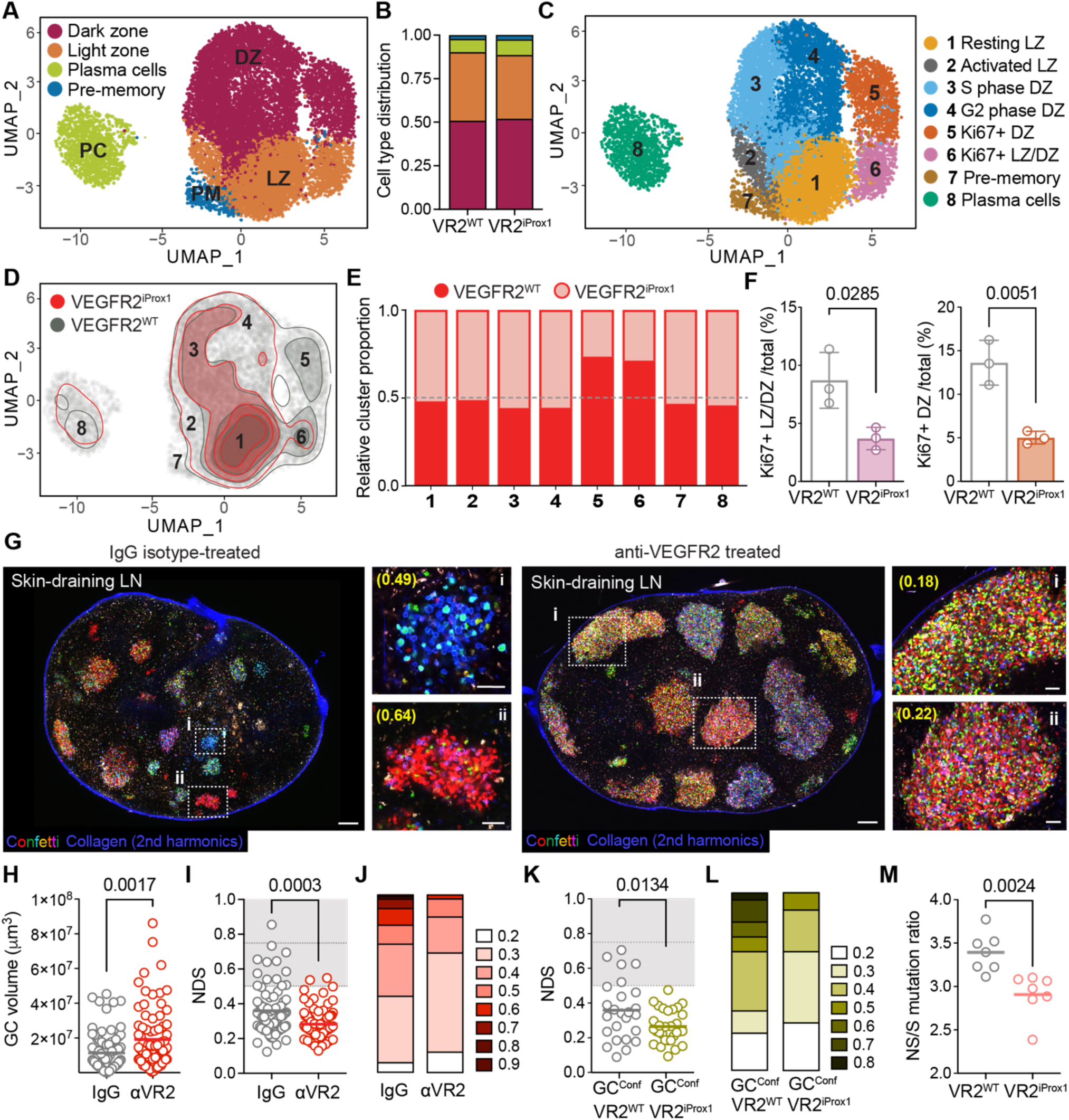
Germinal center selection is impaired in the absence of VEGFR2-dependent lymphatic remodeling. (**A**) UMAP projection of scRNA-seq transcriptomic profiles of germinal center (GC) B cells (B220^+^CD38^−^GL7^+^) and plasma cells (PC, B220^lo^CD138^+^) isolated from draining lymph nodes (dLN) from 3 VEGFR2^WT^ and 3 VEGFR2^iProx1^ mice at day 25 post vaccinia virus (VACV) infection (by scarification). Light zone, LZ; dark zone, DZ. (**B**) Proportional distribution of each cluster in (A) separated by genotype VEGFR2^WT^ (VR2^WT^) and _VEGFR2iProx1 (VR2iProx1)._ (**C**) Higher-resolution unsupervised Seurat clustering of UMAP projection shown in (A), with cell cycle analysis and LZ/DZ annotations, colored by Seurat cluster, numbered and labeled as indicated. (**D**) Contour plot of UMAP in (C) showing cluster density colored by genotype (VEGFR2^WT^, grey; VEGFR2^iProx1^, red). (**E**) Proportional distribution of genotype per cluster (VEGFR2^WT^, red; VEGFR2^iProx1^, light red). Numbers correspond to clusters defined in (C). (**F**) Percent of Ki67^+^ LZ/DZ (cluster 6) (F) and Ki67^+^ DZ (cluster 7) (G) among total sequenced B cells. Each point is one mouse and p values were calculated using two-tailed Student’s t test. (**G**) Representative multiphoton images of dLNs from AID-Confetti mice treated with IgG isotype control (left) or αVEGFR2 blocking antibody (right). (i and ii) indicate GCs shown at higher magnification, with yellow numbers in brackets corresponding to normalized dominance score (NDS). Scale bars = 200 μm (overview) and 50 μm (close-up). (**H**) Quantification of GC volume in AID-Confetti mice treated as in (G). Each point is one GC. (**I**) Quantification of NDS in GCs from (G). Gray shaded area denotes NDS >0.5, dotted lines denote NDS = 0.5 and 0.75. Data are from 2 independent experiments (n=4 mice, IgG; n=5 mice, αVEGFR2). p values were calculated using two-tailed Student’s t test. (**J**) Binned distribution of NDS values for Confetti-labeled GCs in (G). (**K**) Quantification of NDS from VEGFR2^WT^ or VEGFR2^iProx1^ chimeric mice receiving AID-Confetti bone marrow. Each point is one GC. Data are from 3 independent experiments (n=4 mice, AID-Confetti> VEGFR2^WT^; n=3 mice, AID-Confetti> VEGFR2^iProx1^); p values were calculated using two-tailed Student’s t test. (**L**) Binned distribution of NDS values for Confetti-labeled GCs from chimeric mice in (K). (**M**) Ratio of non-silent (NS) to silent (S) BCR mutations in the Ig heavy chain of individually sorted GC B cells (B220^+^CD38^-^GL-7^+^), sequenced via plate-based single-cell V(D)J sequencing. Each point the average per cell NS/S ratio per mouse. Data are from two independent experiments (n=7 mice, 278 cells VEGFR2^WT^; n=7 mice, 299 cells VEGFR2^iProx1^). p values were calculated using two-tailed Student’s *t* test.

To test the hypothesis that GC selection is compromised in the absence of VEGFR2-dependent lymphatic remodeling, we turned to the use of the AID-Confetti multicolor fate-mapping model (*Aicda*^CreERT2/+^.*Rosa26*^Confetti/Confetti^), in which the Brainbow cassette can be conditionally activated by tamoxifen specifically in GC B cells participating in an ongoing GC reaction. In these mice, GC B cells are stochastically colored at peak GC time points, and the emergence of single-color GCs over time indicates successful clonal selection (**Figure S4A**). Importantly, this tool enables the readout of GC function on an individual GC basis, allowing us to observe both inter and intranodal heterogeneity as a function of infection. AID-Confetti mice were infected with VACV as previously described and treated with either isotype control of a VEGFR2-blocking antibody (DC101), which we previously demonstrated to replicate the lymphatic-specific knockout with respect to lymphatic capillary remodeling, viral dissemination, and altered immune surveillance^8^. Antibody treatments were administered one day prior to infection and for every three days until endpoint. Here, consistent with our findings in VEGFR2^iProx1^ mice, anti-VEGFR2–treated AID-Confetti mice showed reduced perifollicular lymphangiogenesis 5 days post-infection compared to isotype control–treated mice (**Figure S4B**). Following infection, mice were treated by oral gavage with tamoxifen to induce Confetti expression at peak GC time points (day 14 and 16) and analyzed 18-20 days later to evaluate the degree of clonal selection **(Figure S4A)**. While in isotype treated controls, we observed the presence of single-color GCs, consistent with clonal selection and emergence of a single, heavily-expanded B cell clone^23,24^, in VEGFR2-antibody treated mice almost all GCs maintained multiple colors (**Figure 3G**) and were significantly larger than controls as measured both by the average radius (**Figure S4C**) and volume (**Figure 3H**). We found no difference in overall number of GCs (**Figure S4D**) or GC turnover (entry and exit dynamics) as measured by colored cell density over time (**Figure S4E**). To quantify observed differences, we used normalized dominance score (NDS)—a measure of clonal dominance within a GC based on color representation, which accounts for unlabeled cells within the GC^24^. An NDS of >0.5 is consistent with clonal selection, whereas an NDS of >0.75 indicates homogenizing selection (a single clone expanding to dominate an entire GC). VEGFR2–antibody treated mice exhibited a marked reduction in NDS compared to controls (**Figure 3I and J**), generally failing to reach an NDS> 0.5. Importantly, these data were validated in bone marrow chimeras, where AID-Confetti bone marrow was transferred into either VEGFR2^WT^ or VEGFR2^iProx1^ mice (**Figure S4F**). In chimeric mice, *Vegfr2* was knocked out prior to infection as in previous experiments. Mice were then infected as described above and labeled with tamoxifen at GC time points **(Figure S4F)**. Tamoxifen-mediated deletion of *Vegfr2* performed prior to infection labelled a proportion of constitutive GCs in the mesenteric LN but induced negligible labeling in VACV-induced GCs in skin-draining LNs prior to re-exposure to tamoxifen. (**Figure S4G and H**). Analysis of NDS at endpoint again demonstrated a significant impairment in the ability of individual GCs to undergo clonal selection (**Figure S4I and 3K and L**). Finally, to provide further functional evidence supporting less effective selection, we sorted single GC B cells from VEGFR2^WT^ and VEGFR2^iProx1^ mice into 96 well plates and performed DNA sequencing of immunoglobulin V(D)J genes. Here, we found that GC B cells in VEGFR2^iProx1^ mice, while still undergoing somatic hypermutation (**Figure S5**), were less likely to select clones with coding mutations that cause amino acid replacements and exert functional impacts on the BCR, and instead had a higher proportion of silent, non-coding mutations (**Figure 3M**). These data together show that the GCs in VEGFR2^iProx1^ mice, while structurally normal, were less able to drive competitive clonal selection and thereby exhibited reduced effectiveness of their systemic antibody repertoire.

### Germinal center size is optimized to promote T-B cell interactions for clonal selection

The emerging picture from our data indicated that the failed humoral immunity observed in mice deficient in VEGFR2-dependent lymphatic remodeling depended upon early spatial regulation of the B cell follicle and subsequent GC volume. We therefore hypothesized that the size of the GC structure could optimize B cell dynamics and clonal selection and in agreement with this, in wild-type mice which exhibit a natural range of GC sizes, GCs that reached an NDS score consistent with clonal selection were almost always significantly smaller than those that did not. We found a negative correlation between NDS and GC size (radius, μm; **Figure 4A**) with GCs larger than 160 μm never reaching color dominance within a labeling window of day 14-20 days post VACV infection (**Figure 4B**). To mechanistically explore the impact of GC volume on selection dynamics we turned to an *in silico* approach using a previously developed model of GC selection^25^. Here, we systematically scaled the number of founder cells within the model to increase the GC volume, consistent with our experimental data (**Figure 2**). We modeled four GC follicle sizes with increasing reaction sphere radii of 160, 208, 240, and 320 μm, which represented the range in GC sizes present in our VACV data (**Figure S4C**). Strikingly, the model predicted a dramatic reduction in clonal dominance (the percent of GC B cells from the same clone per GC; **Figure 4C**) and mean GC B cell affinity (**Figure 4D**) as the GC reaction radius increased from 160 μm to larger radii. The model further predicted that GC B cells in larger GCs, despite the scaling of both FDC and T_FH_ networks, would have reduced *c-Myc* expression (**Figure 4E**), indicating a deficiency in the signaling required to promote B cell selection and regulate extent of DZ proliferation^26,27^. To see if this prediction was corroborated in our experimental data, we turned to the scRNAseq data (**Figure 3**) and indeed observed a significant reduction in *c-Myc* expression, the expression of Myc-target genes, and expression of the related cell-cycle regulator, *Ccnd3,* in LZ populations from VEGFR2^iProx1^ mice relative to controls, particularly in clusters 5 and 6 which were significantly reduced in VEGFR2^iProx1^ mice with associated failures in clonal selection (**Figure 4F and G and S6**).

**Figure 4.**
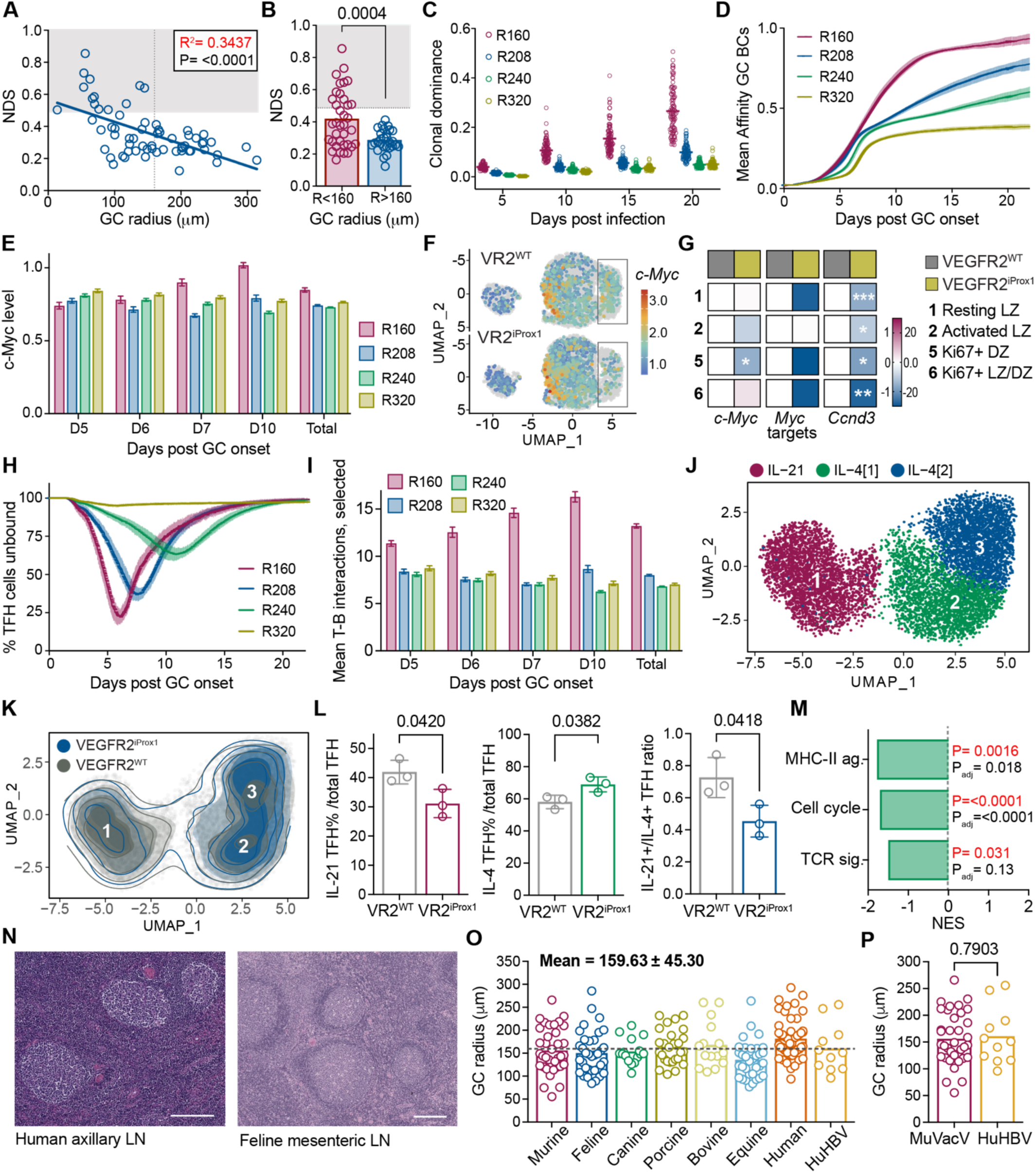
Germinal center size is optimized to promote T cell help and affinity-based clonal selection. (**A**) Correlation between germinal center (GC) radius and normalized dominance score (NDS) in AID-Confetti mice treated with IgG isotype control. Gray shaded area denotes NDS > 0.5, dotted line indicates GC radius = 160 μm. Correlation was determined by simple linear regression with 95% confidence interval. (**B**) NDS values from (A) grouped by GC radius (<160 μm or >160 μm). Each point is one GC. Gray shaded area and dotted line indicates NDS > 0.5. Data in (A–B) are from 2 independent experiments (n = 7 mice). p values were calculated using two-tailed Student’s *t* test. (**C–E**) Computational modeling of GC dynamics across increasing GC radii (radii, mm). Percentage of the dominant clone, *calculated only for GCs with at least 200 cells*, (C), average GC B cell affinity (D), and average cMyc level (E) were modeled as a function of time post-infection or GC onset as indicated. (**F**) UMAP projection of B cell scRNA-seq data from Figure 3, split by genotype (VEGFR2^WT^ (VR2^WT^) control and VEGFR2^iProx1^(VR2^iProx1^)), colored based on the z-scored expression of *c-Myc*. Boxed regions highlight Ki67⁺ DZ and Ki67⁺ LZ/DZ clusters. (**G**) Heatmap showing relative expression (log₂ fold change) of *c-Myc*, *Myc* target genes, and *Ccnd3* across GC B cell clusters, separated by genotype. Asterisks denote significance (\**p* < 0.05, \*\**p* < 0.01, \*\*\**p* < 0.001; Mann-Whitney test). (**H-I**) Computational modeling of follicular helper T cells (T_FH_) –B cell interaction parameters across GC radii as in (C). Percentage of T_FH_ cells not engaged in an interaction with a B cell at a given time (H). Average number of T–B interactions per selected B cell on the indicated day or over the entire duration of the reaction (total) (I). (**J**) UMAP projection of scRNA-seq profiles of sorted T_FH_ cells (TCRβ^+^CD4^+^PD-1^+^CXCR5^+^) from 3 VEGFR2^WT^ and 3 VEGFR2^iProx1^ mice, annotated as IL-21⁺ T_FH_, and two IL-4 expressing T_FH_ populations. (**K**) Contour plot of UMAP in (J), showing density of T_FH_ subsets color coded by genotype (VEGFR2^WT^, gray; VEGFR2^iProx1^, blue). (**L**) Percentage of IL-21⁺ T_FH_ among total T_FH_ (left), IL-4⁺ T_FH_ among total T_FH_ (Middle), and ratio of IL-21⁺ to IL-4⁺ T_FH_ (right) of VEGFR2^WT^ and VEGFR2^iProx1^, quantified from scRNA-seq data in (J). Each point is one mouse; p values were calculated by using two-tailed Student’s *t* test. (**M**) Bar graph showing normalized enrichment scores (NES) for gene signatures related to MHC-II antigen presentation, cell cycle, and TCR signaling in VEGFR2^iProx1^ versus VEGFR2^WT^ T_FH_ cells from scRNA-seq data in (**J**). Significance was determined based on comparing the observed enrichment scores with scores from random permutation testing. Benjamini Hochberg (BH) correction was performed to adjust the p-value to account for multiple testing. (**N**) Representative hematoxylin and eosin (H&E) stained images of human axillary LN (left) and feline mesenteric LNs (right) containing GCs. Scale bars = 200 µm. (**O**) Quantification of GC radius from LN across various mammalian species. Each point is one GC. Dotted line indicates mean GC radius (159.63 µm). (**P**) Comparison of GC radius mouse dLNs at day 15 post VACV infection (MuVACV) and H&E-stained human vaccinated LN biopsies (hepatitis B vaccine, HuHBV). Each point is one GC (n = 3 mice; n = 4 patients). p values were calculated by two-tailed Student’s *t* test.

While *in silico* modeling data paired with scRNAseq analyses identified a deficiency in the B cell signaling required to drive changes in clonal selection, why this would be the case was less clear. Importantly, GC dynamics and B cell clonal expansion depends upon the appropriate and timely acquisition of T cell help in the GC LZ^12,28–30^. While relative frequency of the T_FH_ compartment was preserved in larger GCs, increasing the physical distance of the search space any given B cell would have to traverse to find cognate T cell help, would be expected to lower the efficiency of engaging in successive productive T-B entanglements over the same fixed timeframe. Indeed, when we turned back to the mathematical model, the predicted failure to reach clonal dominance associated with a significant increase in the ‘free’ T_FH_ percentage (T_FH_ cells not actively engaged in an interaction with a B cell) on a population level as a function of GC size (**Figure 4H**), and significantly fewer individual T-B cell interactions expected per selected B cell (**Figure 4I**), indicating a decrease in both the likelihood and magnitude of accumulating timely T cell help. Again, predications from the mathematical model matched our experimental data. Cells sorted from VEGFR2^WT^ and VEGFR2^iProx1^ 25 days post-infection (**Figure S3A**) were subset on cells expressing both *Cd3e* and *Cd4* and re-clustered to reveal seven subclusters (**Figure S7A**). Cluster annotations were defined based on top differentially expressed genes to reveal cytotoxic CD4 (CD4-CTL), cycling CD4, IFNγ^+^ CD4, regulatory T cells (Treg), *Bach2*-expressing CD4, and T_FH_ clusters, either expressing IL-4 or expressing IL-21^31–33^. PCA analysis indicated few significant changes in expression across genotypes within each cluster (**Figure S7B**). However, detailed analysis of the T_FH_ subsets (*Cd4*^+^*Cxcr5*^+^*Pdcd1*^+^*Bcl6*^+^*Icos*^+^; **Figure S7C-E**) defined three T_FH_ clusters, one *Il21*^high^ and two *Il4*^+^. This analysis revealed a significant shift in T_FH_ phenotype, with VEGFR2^iProx1^ mice containing a reduced proportion of *Il21*^high^ expressing T_FH_ compared with *Il21*^low^ negative, *Il4* expressing T_FH_ (**Figure 4J-M and S7C-G**). IL-21 upregulation is strongly associated with T_FH_ differentiation, activation, and maintenance, and is considered to mark active ‘help-givers’ in the GC^33,34^. Indeed, differential expression and pathway analysis revealed a significant reduction in gene programs associated with T_FH_ function, including MHCII antigen presentation, cell cycle and proliferation, and TCR signaling^35^ in T_FH_ taken from VEGFR2^iProx1^ mice relative to controls (**Figure 4M and S7H**). In addition to its feedforward effects on activation of the GC T_FH_ population, IL-21 also acts directly on B cells, augmenting positive selection signals, DZ transition, and increased number of divisions in the DZ^36^. Taken together, these data support a model where GC niche size is an essential determinant of clonal expansion; where oversizing the GC impairs efficiency of T-B engagement, thus reducing positive selection and inertial GC B cell cycling in the DZ^37,38^, which is required to achieve homogenizing selection and GC takeover by selected B cell clones.

Finally, if scaling the GC reaction up past this defined limit of 160 μm truly led to fundamentally inferior performance, we asked whether GC size was maintained in large animals where LNs are significantly larger than those found in mice. To answer this question, we collected archived LN tissues from across a range of mammalian species including cats, dogs, pigs, cows, horse, and humans, and quantified the average GC radius. Consistent with our modeling data that predicted an optimal size for GC function, we found that GCs across all species exhibited a mean radius of 159.63μm (**Figure 4N and O**) with GC number but not individual GC size appearing to scale proportionally in larger LNs. This conservation of GC size was particularly striking when comparing VACV-induced GCs in mouse to human GCs induced following vaccination against hepatitis B virus (**Figure 4P**). These data thereby not only indicate that there is an evolutionarily conserved optimal GC size, but importantly, that the lymphatic vasculature plays an active role in setting these size constraints in the context of VACV infection.

### Viral dissemination to lymph nodes impairs perifollicular lymphangiogenesis and germinal center selection

The implications of these data are that lymphatic remodeling dynamics plays a dual role in optimizing clonal selection during GC formation. Specifically, peripheral lymphatic capillary zippering functions to restrict viral transport to the LN^8^, and perifollicular lymphangiogenesis actively encapsulates activating B cell follicles to create an ideal anatomical niche for GC selection. What remained unclear, however, is whether this selection phenotype could be simply explained by the presence of virus in the draining LN that is observed in VEGFR2^iProx1^ mice^8^. We therefore last sought to understand the extent to which virus itself was sufficient for the observed phenotype and to uncouple the relative effects of peripheral and LN lymphatic remodeling.

To first understand the impact of intact virus on the expansion of B cell follicle and GC response we directly compared mice infected by scarification to a cohort infected by intradermal injection of VACV at the same dose. While scarification induces zippering of peripheral lymphatic capillaries, which restrains viral transport to the LN, intradermal injection rapidly forces virus into naïve lymphatic capillaries through high interstitial fluid pressures, leading to direct infection of the LN^8^. No differences in peak skin titers are observed when comparing infections by these two routes^8^. Five days following infection, while LNs draining skin infected by scarification demonstrated perifollicular lymphangiogenesis as previously demonstrated (**Figure 1**), LNs draining skin infected by intradermal injection demonstrated a reduction in new lymphatic growth, specifically a loss of perifollicular lymphatic structures (**Figure 5A-B**) and an increase in B cell follicle area (**Figure 5C**), consistent with the phenotype observed in VEGFR2^iProx1^ mice. These early changes in lymphatic and B cell dynamics consistently translated to a loss of GC efficiency. When evaluating GC time points in AID-Confetti mice (**Figure 5D**), GCs induced by intradermal infections were significantly larger in volume (**Figure 5E**) and impaired in their ability to select relative to GCs forming in LNs draining skin infected by scarification (**Figure 5F and G**). These data indicate that the presence of virus in the LN, regulated by VEGFR2-dependent peripheral lymphatic zippering, is central to the observed phenotype and could suggest that perifollicular lymphangiogenesis is a readout of productive B cell activation rather than an active player in constraining or optimizing the response.

**Figure 5.**
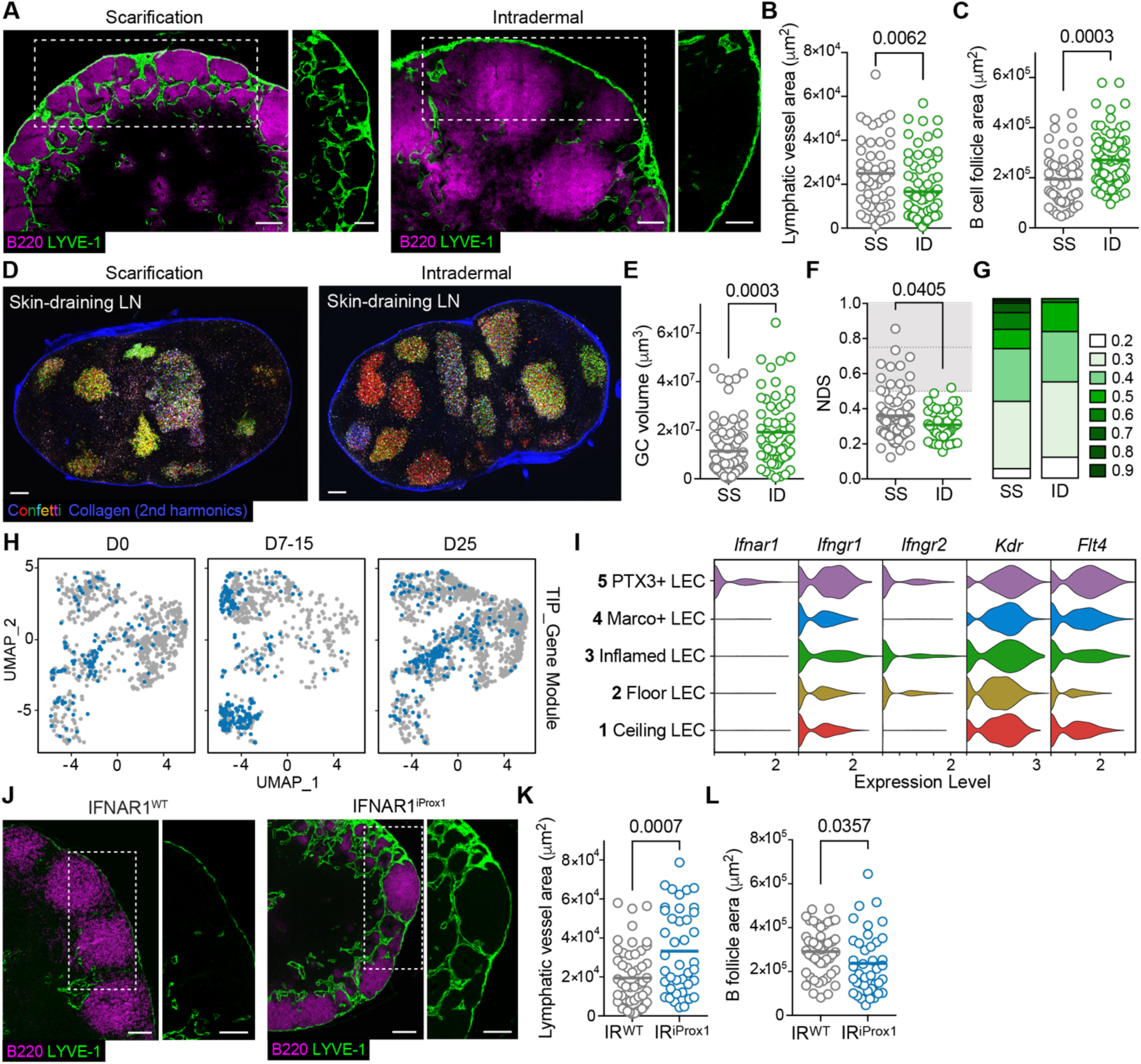
Viral dissemination to lymph nodes reduces germinal center efficiency by inhibiting perifollicular lymphangiogenesis. (**A**) Representative immunofluorescence images of draining lymph nodes (dLN) from C57BL/6 wildtype mice at day 5 post-vaccinia virus (VACV) infection by scarification (SS) or by intradermal (ID) injection. White box indicates high magnification inset. Green, LYVE-1; magenta, B220. Scale bars = 200 μm. (**B**) Quantification of perifollicular lymphatic vessel area in the dLNs from (A). Each point is a follicle. p-values were calculated using two-tailed Student’s t-test. Data in (B-C) from two independent experiments (n=6 mice, SS; n=8 mice, ID) (**C**) Quantification of B follicle area from (A). Each point is a follicle. p-values were calculated using two-tailed Student’s t-test. (**D**) Representative of multiphoton images of dLNs from AID-Confetti mice following VACV infection by SS or ID injection. Scale bars = 200 μm. (**E**) Quantification of germinal center (GC) volume in AID-Confetti mice following VACV infection by SS or ID injection. Each point is one GC. p-values were calculated using two-tailed Student’s t-test. Data are from 2 independent experiments (n=4 mice, SS; n=4 mice, ID). (**F**) Quantification of normalized dominance score (NDS) in GCs from (D). Gray shaded area denotes NDS >0.5, dotted lines denote NDS = 0.5 and 0.75. (**G**) Binned distribution of NDS values for Confetti-labeled GCs in (F). (**H**) Feature plot of LN LECs over time post VACV SS, colored by expression of a ‘tip-like’ lymphatic vessel gene module. (**I**) Violin plots showing expression of indicated genes across LEC clusters. (**J**) Representative immunofluorescence images of dLNs at day 5 post VACV ID injection from IFNAR^WT^ (IR^WT^) and IFNAR^iProx1^ (IR^iProx1^) mice. White box indicates high magnification inset. Green, LYVE-1; magenta, B220. Scale bars = 200 μm. (**K-L**) Quantification of perifollicular lymphatic vessel area (K) B cell follicle area (L) in the dLNs of IFNAR^WT^ and IFNAR^iProx1^ from (J). Each point is one follicle. P values were calculated using two-tailed Student’s t-test. Data presented in (K-L) were obtained from the same experiments, independently repeated twice (*n* = 6 mice, IFNAR^WT^; *n* = 7 mice, IFNAR^iProx1^).

Lymphangiogenesis, however, while driven by growth factor signaling (e.g. VEGFR2 and VEGFR3) is also sensitive to negative regulation in inflammatory microenvironments. Both type I and II IFNs are potent inhibitors of LN^39,40^ and tumor-associated lymphangiogenesis^41^. We recently found that in the absence of IFN-dependent control, proliferating LECs acquired a highly invasive transcriptional state^41^ also seen in a model of lymphatic malformations in skin^42^. This transcriptional state is defined by high expression of *Ptx3*, *Mrc1*, *Ccl2*, and *Ackr2* and a module of genes associated with a tip-like cell state. It was therefore notable that the LEC population that significantly expanded in VACV-draining LNs following infection by scarification was the PTX3+ LEC subset, which shares a similar transcriptional program (**Figure 1**). We therefore projected a module score based on a tip-like signature that is enriched in proliferative and invasive tumor-associated LECs^41^, and found that this signature was particularly elevated in the PTX3 parent and inflamed populations at days 7-15 post infection (**Figure 5H**), time points consistent with LEC expansion and GC activation. Furthermore, while all LEC subsets exhibited relatively similar levels of expression of pro-growth receptors, *Kdr* and *Flt4* (VEGFR2 and VEGFR3, respectively), there was variable expression of the type I interferon receptor (*Ifnar1*) and the beta chain of the type II IFN receptor (*Ifngr2*) transcripts, with only the PTX3+ LEC subset appearing to be poised for response to both types of IFN in the LN (**Figure 5I**). Based on these data, we hypothesized that perifollicular lymphangiogenesis could be regulated by the presence of virus in the LN by way of sensing an altered inflammatory environment. Therefore, rather than the loss of lymphangiogenesis seen in VEGFR2^iProx1^ mice being driven by the absence of a pro-growth signal, the failure to proliferate could be a consequence of local virus-induced IFNs that restrained growth *in situ*. This conceptual shift was further supported by the fact that despite the high levels of VEGFA expression seen in B cells activated following immunization with the model antigen keyhole limpet hemocyanin emulsified in complete Freund’s adjuvant^16^, little to no *Vegfa* transcript was observed in activated B cell follicles and GC B cells following VACV infection in wildtype mice (data not shown). We therefore postulated that if we could protect LN lymphatic vessels from an inhibitory, anti-viral IFN signal, we could determine if perifollicular lymphangiogenesis was sufficient to rescue GC control– even in the presence of abundant LN virus. To test this hypothesis, we repeated intradermal VACV infections in mice with lymphatic-specific loss of the type I IFN receptor (*Ifnar1*; IFNAR1^iProx1^). As before, *Ifnar1* deletion was induced prior to infection and the lymphatic and B cell response was quantified 5 days post-infection. Although again, intradermal infection failed to trigger perifollicular lymphatic growth in IFNAR1^WT^ controls, in mice whose LN lymphatic vessels were insensitive to type I IFNs (IFNAR1^iProx1^) we saw robust perifollicular lymphangiogenesis (**Figure 5J and K**), and a striking rescue of B cell follicle constraint (**Figure 5L**), creating follicular structures phenocopying those observed in the context of wildtype infection by scarification. Collectively, these data indicate that lymphatic vessels directly optimize GC dynamics, first by regulating the flow of virus and viral antigens to the LN, which in turn preserves a LN microenvironment favorable to the induction of a perifollicular lymphangiogenic response. Perifollicular lymphangiogenesis then actively constrains GC size to facilitate efficient positive selection by T cells and optimal clonal expansion.

## Discussion

Despite the critical role of functional lymphatic transport for the delivery of antigen to draining LNs, whether and how lymphatic vessels, themselves, actively regulate transport and thereby adaptive immune responses remain poorly understood. We previously made the surprising finding that lymphatic vessels actively constrain tissue efflux following infection by scarification, serving to sequester virus at the site of infection while simultaneously optimizing antigen presentation and CD8 T cell priming. Here we build on that finding to investigate the impact of viral-induced lymphatic remodeling events on the humoral response to infection. We demonstrate that active, VEGFR2-dependent remodeling of the lymphatic vasculature, both in the periphery and in the draining LN, is required to optimize GC B cell selection and support the production of high titers of protective virus-specific antibodies. Importantly, the lymphatic vasculature optimizes this response first by constraining viral transport to the draining LN, which establishes a local inflammatory context that is permissive to rapid LN lymphangiogenesis. This LN lymphatic growth encapsulates activating B cell follicles and constrains their expansion, generating GC niches right-sized for search dynamics conducive to competitive T_FH_-meditated selection. Importantly, we find that this optimal GC size is conserved across mammalian species, indicating that size is as a key determinant of successful B cell selection, and therefore designating a fundamental role for the lymphatic vasculature in setting these critical size constraints in the context of VACV infection. These data therefore support an emerging understanding that the lymphatic vasculature is an active, structural scaffold that serves to compartmentalize antigenic and inflammatory signals to shape the adaptive immune response following peripheral tissue challenge.

Our model places lymphatic transport to the LN as the first key factor determining the footprint of the B cell follicle during activation, which in turn, regulates downstream GC dynamics. At steady state, antigen transport to the LN is largely size dependent, with first peripheral lymphatic capillaries and then SCS LECs and LN conduits, constituting a physical filtration system that restricts large, complexed antigens to the LN periphery and allows low molecular weight antigens direct access to conduits. B cell activation depends upon the handoff of antigen to FDC networks thought to be mediated by the direct scavenging of lymph-borne pathogens or antigen complexes by the SCS macrophage^43^. Rather than being static, however, the size exclusion properties of the lymphatic system change as a function of environmental cues^8,44^, increasing barriers to free, paracellular antigen transport and requiring more active mechanisms of delivery to the LN. Therefore, how antigen delivery to naïve B cells is accomplished in the context of VACV scarification remains an open question, but with several emerging possibilities. First, peripheral lymphatic zippering may not be a complete block on antigen transport, as it takes at least 12 hours to begin to see peripheral lymphatic zippering^8^, which may leave enough time for early, rapid priming. Importantly, however, LECs are capable of dynamin-dependent vesicular transcytosis that mediates the transfer of large protein antigens, including lymph-borne antibodies^45^, and is enhanced by inflammatory fluid flows^46^, indicating that LECs may switch to active, selective antigen transport mechanisms. Finally, DCs can capture and re-expose viral particles^47^ and immune complexes^2^ raising the possibility of active, cell-based antigen transport from the periphery for presentation to B cells^48,49^. Antigen delivered by APCs may more efficiently activate naïve B cells^50^, and favor FDC loading^50,51^. Interactions with membrane-complexed antigen are particularly important in the GC^52^, where GC B cells are largely agnostic to soluble antigen stimulation^52,53^ and instead use mechanical force to regulate BCR signal strength and antigen acquisition from FDCs during affinity discrimination^52,54^. A shift from passive to more active mechanisms of transport are likely to delay the delivery of antigen to FDCs over time relative to bolus injection. This natural ‘low and slow’ delivery system may specifically support long-lasting, more effective GC responses as also seen when vaccines are dosed slowly over time^3,55–57^.

The generation of high-affinity antibodies from GCs revolves around expanding B cells most successful in competing for two main limiting resources: antigen and T cell help. Despite GCs lacking lymphatic constraint becoming oversized, they appeared to be otherwise phenotypically normal in terms of FDC and T_FH_ zonal distribution, the inherent ability to mutate their V(D)J genes, and transcriptional profile of GC T and B cell subsets all arguing against any direct effects of LEC signaling on lymphocytes. Instead, we propose that GC niche size alone-albeit inherently set by lymphangiogenesis-is sufficient to account for the observed changes in GC clonal selection. This expansion in GC size seems to impact selection by limiting the accumulation of T cell help over time. Predicted by mathematical modeling, this was substantiated directly in our transcriptional analysis of both GC B and T cells. In VEGFR2^iProx1^ mice, GC B cells dropped out at the transition from LZ to DZ, consistent with less recent positive selection via T cell help and impaired induction of *Myc* and *ccnd3* expression, which directs the signal-free inertial cell cycling required to complete a clonal burst in the DZ^37^. Similarly, GC T cells exhibited signs of reduced activation, where a downregulation of TCR signaling and cell cycle gene pathways in VEGFR2^iProx1^ mice are consistent with reduced antigen engagement, activation, and proliferation within the T_FH_ population that occurs following GC B cell interaction^35^. We also identify a specific reduction in *Il21*^+^ T_FH_ cells in relation to *Il4*^+^ cells. Although the full implications of this shift remain unknown, IL-21^+^ T_FH_ are known to play profound roles in the earlier phases of the GC response to regulate B cell selection, proliferation, cyclic re-entry and clonal expansion, whereas IL-4^+^ T_FH_ are more associated with plasma cell fate decisions^20,31,32,34^. Taken all together, inefficient T-B cell interactions are likely to limit the ability of individual GC B cell clones to gain a competitive advantage over their neighbors. In accordance with these data, we show that clonal bursts were absent in VEGFR2^iProx1^ GCs, which was accompanied by less affinity maturation (evidenced by reduced selection of productive mutations) and less protective antibody production.

Although our findings position GC niche size as a previously unappreciated rheostat of GC efficiency, how the perifollicular lymphatic response achieves this GC size regulation remains unresolved. Our data demonstrate that the release of virus into the LN prevents a lymphangiogenic response through the local production of type I IFNs, such that growth and GC constraint can be rescued through lymphatic-specific deletion of the IFNAR1. Therefore, the effect of LN virus is in part indirect, exerting its effect on GC size through the establishment of a bounding perimeter. And so, the establishment of GC niche area could simply be a consequence of a physical ‘fencing’ of the B cell follicle, or alternatively be more actively controlled by lymphatic vessels via secretion or sequestration of chemokines that maintain lymphocyte migration gradients^58,59^. Interestingly, the LEC subset that likely composes this perifollicular boundary, PTX3^+^ LECs, exhibits a unique transcriptional profile that may suggest active mechanisms of B cell engagement. It is intriguing to note that PTX3, itself, when released by neutrophils binds splenic marginal zone B cells and promotes IgM and IgG production after infection^60^; that as an fibroblast growth factor (FGF)-trap, PTX3 may regulate local proliferative potential^61^; and that PTX3 appears to sequester cell remnants away from APCs^62^, which may be required to limit autoimmunity.

Finally, the initiating signal that drives perifollicular lymphangiogenesis remains uncertain. B cells may actively shape the LN lymphatic network, and at least in one vaccine setting, produce VEGFA and induce LN lymphangiogenesis^16^. Consistent with this, either the transgenic expression of VEGFA in B cells (CD19-Cre), or enhancing the B cell compartment (Em-c-Myc mice)^63^, results in expanded lymphatic vessel networks. Interestingly, immune complexes are also sufficient to induce LN lymphangiogenesis through induction of VEGFA in SCS macrophages and dendritic cells^64^, which may suggest that early antigen capture initiates concordant lymphatic growth. While this direct association between B cells, VEGFA, and lymphatic vessels may be operational in our system, it is intriguing that we see very little VEGFA in VACV dLN and the loss of IFN signaling is sufficient to rescue lymphatic growth in the LN, suggesting at least that a balance between pro- and anti-growth signals is critical to shaping lymphatic structures^39–41^ and GC responses. These data imply then that lymphatic dynamics may tune the abundance and breadth of systemic humoral responses. We identified significant size variation across the natural LN GC response, and accordingly, achieving monoclonality in the GC is somewhat of a ‘jackpot’ event. This is true across many settings, including viral infections, immunizations, autoimmunity, and in naturally occurring mucosal GCs^23,24,65,66^. Whilst extensive clonal selection evolves very high antibody specificity and affinity- of utmost importance for rapid pathogen clearance during infections-there is significant benefit for both the primary and memory responses to evolve a combination of binding specificity and breadth to ensure complete coverage of the antigen landscape. The aberrant generation of antibodies is not only detrimental during infections or vaccination, but is associated with immune related adverse events and autoimmunity^67,68^.

In conclusion, our data define a fundamental size limitation to productive GC responses that is conserved across mammalian species despite increasing LN size. We find an increasing number of size-regulated GC niches in the LNs of larger mammals, rather than a fixed number of GC structures that increase in volume with LN size. A central regulator of GC size is the lymphatic vasculature, which serves firstly to limit viral dissemination to the LN and protect the tissue from aberrant inflammatory signals, and secondly encapsulates the growing B cell follicle to set inter-follicular boundaries, limit follicle expansion and thereby optimize the efficiency of competitive clonal selection. While the work here was performed using VACV, which is a relatively large, complex, enveloped virus, it remains to be seen how other viruses and bacteria may bypass these lymphatic-dependent mechanisms of constraint and the impact this exerts on the fitness of GC responses. We therefore propose that the active mechanisms that govern lymphatic-dependent transport and compartmentalization may be important innate determinants of the host-pathogen response and present new strategies to regulate vaccine biodistribution, an important design parameter for novel vaccine development.

## Resource Availability

Further information and requests for resources and reagents should be directed to and will be fulfilled by the lead contacts.

## Data and code availability

Sequencing data will be made publicly available upon acceptance. The full code, parameter files and documentation for the mathematical modeling are available on Zenodo^69^.

## Acknowledgements

The authors acknowledge computational biology support from Igor Dolgalev and Ai C. Ra at the Translational Immunology Center (TrIC) at NYULH, and Tiago B. R. Castro at the Laboratory of Lymphocyte Dynamics at the Rockefeller University. Authors extend their thanks to the histology staff at the New York State Animal Health Diagnostic Center for case processing and slide scanning. Authors Marta Schips and Niklas Schwan contributed equally.

## Contributions

Conceptualization: CRN and AWL

Methodology: TM, GB, KZ, MS, NS, VC, WF, MC

Investigation: TM, GB, KZ, MS, NS, LFC, OI, LF, SS, HC

Funding acquisition: CRN and AWL Specimens: RH, ADM

Supervision: RH, SK, MM-H, CRN, and AWL

Writing – original draft: CRN, AWL

Writing – review & editing: TM, GB, KZ, MS, NS, LFC, OI, VC, WF, LF, SS, HC, MC, RH, SK, ADM, MM-H, CRN, AWL

## Methods

### Mice, Housing, and Animal Procedures

All animal procedures were approved and performed in accordance with institutional guidelines for the care and use of laboratory animals at the New York University Langone Health. Mice were maintained under specific pathogen–free conditions in facilities at NYU, with a 12-hour light/12-hour dark cycle at a constant temperature (20–23 °C) and given ad libitum access to normal chow diet and water.

The following mouse strains were used: C57BL/6J (stock no. 000664), *Ifnar1*^fl/fl^ (stock no.028256), and *Vegfr2*^fl/fl^ (stock no. 018977) were purchased from The Jackson Laboratory. Prox-1:CreERT2 mice^70^ were provided by V.H. Engelhard (University of Virginia, Charlottesville, VA) in agreement with T. Makinen (Uppsala University, Uppsala, Sweden) and bred to the indicated floxed alleles in house. *Aicda*^CreERT2^ (*Aicda*^tm1.1(cre/ERT2)Crey^)^71^ and *Rosa26*^Confetti^ ^72^ (collectively referred to as AID-Confetti) were provided by G. Victoria (Rockefeller University, New York, NY) and bred in house. For Prox1CreERT2 mice, Cre recombination was induced by administering 2 mg tamoxifen (Sigma T5648) dissolved in corn oil (Sigma, C8267) at 20 mg ml^−1^ via intraperitoneal injection (i.p.) for five consecutive days, followed by a one-week rest period before experiments. For AID-confetti mice, Cre recombination was induced by two gavages of 10 mg tamoxifen dissolved in corn oil at 50 mg ml^−1^, 2 days apart.

VEGFR2 blocking antibody (InVivoMAb DC101, Bio X Cell, A250834; 200 µg) was administered intraperitoneally on days 0 and 3. BrdU (Fisher Scientific, H2726006) 2.0 mg/mouse was injected i.p. on the day of infection and provided in drinking water (0.8 mg/ml) for five days. For in vivo experiments, age- and sex-matched mice (8–14 weeks old) were used, with at least 3 mice per group. Each experiment was independently repeated at least twice. Littermate controls, tamoxifen-induced controls, or isotype antibody (InVivoMAB anti IgG1, BE0088) controls were used in all experimental designs.

### VACV expansion and infection

Vaccinia virus (VACV) was propagated in BSC-40 cells using standard procedures^73,74^. For cutaneous infection by scarification, mice were inoculated on the dorsal ear pinna by 25 punctures with a 29-gauge needle following topical application of 5 × 10⁶ PFU VACV in 10 μl PBS. For intradermal infection, 5 × 10⁶ PFU VACV in 10 μl PBS was injected into the ear tip using a Hamilton syringe.

### Immunofluorescence microscopy

Tissues were fixed in 1% paraformaldehyde (Thermo Scientific, AAJ19943K2) for 24 hours at 4°C and transferred to 15% sucrose (Sigma, S9378) (overnight at 4°C) followed by 30% sucrose (overnight at 4°C). Tissues were then embedded in O.C.T. Compound (Fisher Scientific, 23-730-571) and frozen in 2-methylbutane (Sigma, 320404) in liquid nitrogen. Whole lymph nodes were cryosectioned at 10–30 µm thickness using a Leica Cryotome, and sections containing GCs at comparable depths from the LN surface were selected for staining. Tissue sections were blocked and permeabilized using 2.5% BSA (Fisher Scientific BP1600), 0.1% Triton X (Sigma, X100), and 0.3 M glycine (Sigma, G8898) in PBS for 30 minutes at RT. Sections were stained with primary antibodies in blocking buffer overnight at 4C. Primary antibodies were purchased from ReliaTech, eBioscience, BioLegend, and BD Biosciences: LYVE-1 (103-PA50S), Ki-67 (SolA15), B220 (RA3-6B2), CD3e (145 2C11), CD19 (6D5), CD35 (8C12).

Slides were washed with PBS two times before stained with secondary antibodies in blocking buffer for 30 minutes at RT. Secondary antibodies were purchased from Invitrogen: anti-Rabbit IgG (H+L) AF647, anti-Rat IgG (H+L) AF647, anti-Rat IgG (H+L) AF546, anti-hamster IgG AF488, and Streptavidin AF546 (Invitrogen). After three washes, slides were mounted with Prolong Diamond Antifade (Invitrogen, P36961) or Vectashield (Vector Laboratories, H-1700) mounting media on 1.5 mm coverslips and imaged on Keyence BX-X810 microscope after curing.

### Hematoxylin and eosin (H&E) staining

A sub-cohort of four healthy adult males (ages 35–50) enrolled in a Hepatitis B (HepB) vaccine study at NYU Langone Health (NCT04674462) was analyzed for germinal center size. Two participants with prior HepB vaccination received a single booster dose, while two vaccine-naïve participants received a standard three-dose vaccine series administered at approximately 0, 1, and 6 months after the initial visit. Core needle biopsies were performed 10–21 days after booster or final vaccine dose. Ultrasound-guided biopsies of ipsilateral axillary lymph nodes were performed with an 18-gauge needle by NYU Langone Interventional Radiology. Biopsied cores were immediately placed in RPMI and transported to the NYU Center for Biospecimen Research and Development (CBRD). One core from each participant was formalin-fixed and paraffin-embedded for histologic analysis.

A retrospective search was conducted in the archives of the NYU CBRD (RRID:SCR_017930) for human lymph nodes. A similar search was performed from the Section of Anatomic Pathology, New York State Animal Health Diagnostic Center, Cornell University College of Veterinary Medicine for LNs from other non-human, mammalian samples. The keywords “lymph node” and “hyperplastic” were used, yielding three unique cases each from canine, feline, bovine, equine, and porcine samples. All tissues were processed using standard histologic H&E staining protocols, performed by the NYU Langone Experimental Pathology Core for human samples and the Cornell University College of Veterinary Medicine for mammalian samples. Slides were digitally scanned using a Leica AT2 scanner (human samples) or a Ventana DP200 slide scanner (mammalian samples) and analyzed for the presence of germinal centers.

### Image analysis

Perifollicular lymphatic vessel (LYVE-1) area was measured by an ImageJ macro pipeline. First, B follicle masks were manually drawn. From there, the macro generates a band with a total 180-micron thickness outside of the drawn B follicle mask and measures LYVE1+ area within the band (perifollicular lymphatic vessel area).

GC diameter from different mammalian species were measured using Animal Health Diagnostic Center Annotation Tools (Cornell University) or ImageJ software. The GC diameter was measured across two perpendicular axes and averaged.

### Quantification of selection in AID-Confetti mice

Color dominance in AID-Confetti GCs was determined in 3D datasets reconstructed using ImageJ software and the Bio-formats plugin. Cells of each color or color combination were counted manually in two or more Z-planes, at least 20μm apart, using the Cell Counter plugin on ImageJ. The normalized dominance score (NDS), a measure of how single-colored a given GC is considering both colored and uncolored cells, was calculated using previously published parameters^23^. Briefly, for each GC, we calculated the proportion of Confetti-labeled cells by determining the density of fluorescent cells per 100 μm² in anatomically defined DZ regions where cell distribution is homogeneous. The proportion of labeled cells was then multiplied by the fraction of cells expressing the dominant Confetti color combination to establish the true dominance of that color within the entire GC. Fully recombined GCs have a cell density very close to 1, making an NDS of 1 a good approximation of 100% GC occupancy by the dominant color.

### Multiphoton imaging

Superficial cervical dLNs were excised and prepared for imaging as described previously^23^. Briefly, adipose tissue was removed under a dissecting light microscope and LNs were mounted in PBS between two coverslips sealed with vacuum grease. Imaging was performed on an Olympus multiphoton FVMPE-RS with a MaiTai DeepSee tunable Ti:Sapphire laser (Spectra-Physics) set to 930 nm, a 25X/1.05 XLPLN25XWM objective lens with zoom set to a pixel size of 0.8 um, narrow band blue/green/red bandpass filters, and FluoView software.

### VACV plaque assay

Ears and lymph nodes (LNs) were homogenized in 0.5–1 ml RPMI 1640 (HyClone SH30027.FS) supplemented with 1% heat-inactivated FBS using a handheld tissue homogenizer. Homogenates underwent three freeze–thaw cycles with vortexing, followed by serial dilution. Diluted samples were applied to BSC-40 cell monolayers and incubated for 48–72 h under an agarose gel overlay. After overlay gel removal, plaques were visualized by counterstaining with 1% crystal violet (Sigma, C6158) in PBS containing 2% ethanol, and plaque-forming units (PFU) were quantified as previously described^75^.

### Flow cytometry

To generate single cell suspensions for lymphocyte analysis, LNs were mechanically dissociated using disposable micropestles (Axygen, PES-15-B-SI) in 100 μL PBS supplemented with 0.5% BSA and 1 mM EDTA (Sigma, EDS-500G) (FACS buffer). For analysis of lymph node stromal cells (LNSC), pooled LNs were minced with surgical scissors and digested sequentially in digestion medium (I) containing collagenase IV (1 mg/mL; Worthington Biochemical, LS004186) and DNase I (>80 U/mL; Roche, 04536282001), followed by digestion in medium (II) containing collagenase D (1 mg/mL; Sigma, 11088866001) and DNase I (>50 U/mL). In all cases, single cell suspensions were filtered through 70-μm strainers, stained with antibody panels diluted in FACS buffer supplemented with Fc-block for 20 min at 4°C, and fixed in 1% paraformaldehyde (PFA) at 4°C overnight prior to flow cytometry analysis. Antibodies were purchased from BioLegend, eBioscience, BD Biosciences, and Tonbo and included: Ghost Dye Red 780 and BV510 for dead cell exclusion, CD8 (53-6.7), CD45 (104), CD4 (GK1.5), VEGFR2 (AVAS12), LYVE-1 (ALY7), CD274 (MIH5), TCR α/β (H57-597), CD3 (17A2), CD138 (281-2), GL7 (GL7), CD31 (MEC13.3), gp38 (8.1.1), CD38 (HB-7 or 90), PD-1(29F.1A12), CXCR4 (12G5), B220 (RA3-6B2), Fas (Jo2).

Intracellular BrdU staining was performed following surface staining of single-cell suspensions, after which cells were fixed in BD Cytofix/Cytoperm buffer (BD Biosciences, 554722) on ice for 15–30 minutes. Fixed cells were washed in BD perm/wash buffer (BD Biosciences 554723) and incubated on ice in BD Cytoperm Plus buffer (BD Biosciences 561651) for 10 min. Cells were refixed for 5 min and treated with DNAse (300 μg/ml) for 1.5 h at 37°C and then stained with anti-BrdU (BioLegend Bu20a FITC, 1:25) for 20 min at RT. Data were acquired on a BD Biosciences Symphony A5 flow cytometer and analyzed using FlowJo software (Treestar Inc.).

### ELISA

Total immunoglobulin (Ig) levels in serum from each mouse were quantified and normalized to ensure equal loading for detection of VACV–specific antibodies. Serum Ig concentrations were measured by ELISA using high-binding microplates (Corning, 3690) coated with Goat anti-mouse Ig (1:1000, SouthernBiotech, 1010-01). Serum samples were serially diluted four times, starting at 1:500 with a 4X dilution factor. A standard curve was generated on each plate using serial dilutions of purified total Ig standards, starting at 200 ng/mL and serially diluted eight times starting at 2X dilution factor. Isotype-specific detection antibodies were used to detect IgG (Invitrogen, cat # 62-6520), IgM (Invitrogen, cat# PA1-84383), IgG2c (Southern Biotech, cat# 1077-05) and IgG2b (Southern Biotech, cat# 1090-05). Sample Ig concentrations were interpolated from the standard curve, and equal amounts of total Ig from each sample were used for subsequent VACV-specific ELISAs. Vaccinia virus–specific ELISAs were then performed using high-binding 96-well plates coated with inactivated lysates from VACV (WR strain)–infected BSC-40 cells (1:800 dilution). Plates were blocked with 1% BSA in PBS for 1 h at room temperature, then incubated overnight at 4 °C with equal amounts of total Ig from each serum sample. Bound antibodies were detected using horseradish peroxidase (HRP)–conjugated goat anti-mouse IgG (Jackson ImmunoResearch, 115-035-003), IgM (115-035-020), IgG2c (115-007-188) +IgG2b (115-007-187). In all assays, ELISAs were developed with TMB (slow kinetic form, Sigma). Optical density was measured at 450 nm with a Flexstation 3 Multi-Mode Microplate Reader.

### Bone marrow chimera generation

8–10-weeks old recipient mice received two doses of whole-body radiation (2 x 450 rads, at least 4 h apart) using an x-ray irradiator. Bone marrow was isolated from hind limbs of AID-Confetti mice and 5–10 × 10^6^ cells injected retro-orbitally into recipients. Mice were maintained on 2 mg/ml Ampicillin (Cayman Chemical Company, 14417) antibiotic water changed twice per week for 6 weeks. Mice were rested for a further two weeks without antibiotics prior to performing experimental procedures.

### Single cell RNA-sequencing

For endothelial cell (EC) isolation, cervical lymph nodes from 10 naïve female C57BL/6J mice and draining LNs from 5 VACV-infected female C57BL/6J mice were pooled separately. The tissues were then processed into single-cell suspensions as described in the “Flow cytometry” section. Cells were stained with anti-CD31 APC, anti-PDPN PE-Cy7, anti-CD45 PE, and DAPI for live/dead discrimination. Endothelial cells (CD31^+^) were enriched via positive selection using anti-APC Microbeads (Miltenyi Biotec, 130-090-855) and passed through LS magnetic columns (Miltenyi Biotec, 130-042-401) following the manufacturer’s protocol.

For GC B cells, PCs, and T_FH_ cell isolation, single-cell suspensions from 3 VEGFR2^WT^ and 3 VEGFR2^iProx1^ dLNs were processed as separate samples as described in the “Flow Cytometry” section. Prior to sorting, lymphocytes were stained with anti-TCRβ APC-Cy7, anti-CD138 BV650, anti-B220 BUV395, anti-GL7 FITC, anti-CD38 BUV737, anti-CD4 PE, and anti–PD-1 BV785 (supplementary Figure 3). Each mouse sample was further labeled with antibody hashtag oligonucleotide TotalSeq C (BioLegend).

Stained and enriched cells were sorted into RPMI supplemented with 2% FBS using a FACS Aria III (BD Biosciences; 100 μm nozzle for EC or 70 μm nozzle for lymphocytes; <3,000 cells/s) following sorting strategy: ECs (CD45^-^CD31^+^), GC B cells (TCRβ^−^ B220^+^ GL7^+^ CD38^−^), PCs (TCRβ^−^ B220^mid-lo^ CD138^+^), and T_FH_ cells (TCRβ^+^ CD4^+^ PD-1^+^ CXCR5^+^) (see Supplemental Figure 3). Sorted cells were pelleted, concentrated, and immediately loaded onto the 10x Genomics Chromium platform to generate single-cell gel beads in emulsions. For EC, libraries were prepared using the Chromium Next GEM Single Cell 3′ v3.1 Reagent Kit (16 rxns, PN-1000121; 10x Genomics) according to the manufacturer’s instructions and sequenced on a NovaSeq 6000 (Illumina). For lymphocytes, libraries were prepared using the Single-cell 5’ Gene Expression kit (10X Genomics) according to the manufacturer’s instructions and sequenced on a NovaSeq X Plus (Illumina). Pooled VEGFR2^WT^ samples (29,750 cells) and VEGFR2^iProx1^ samples (50,400 cells) were loaded into separate lanes. Library preparation and scRNA-sequencing were performed at the NYU Langone Health Genome Technology Center. The demultiplexing, barcoded processing, gene counting, and aggregation were made using the Cell Ranger software (v9) and aligned to the mouse genome (mm10).

### Single-Cell RNA-seq analysis

Analysis of single-cell RNA sequencing was performed at NYULMC HPC UltraViolet. Data were processed and visualized using the Seurat (v5.3.0) package in R ((v4.4.2 LEC data sets) and (R4.4.0 lymphocytes data sets)). Quality control filtering was applied to exclude cells with fewer than 100 or more than 10,000 detected genes, >10% (LECs) >5% (lymphocytes) mitochondrial reads, >20% ribosomal protein genes (lymphocytes only), or >25% ribosomal RNA genes. EC datasets from multiple time points post VACV infection were integrated using the Harmony package^76^. The optimal number of principal components was determined using the ElbowPlot function. Cells were clustered using shared nearest-neighbor (SNN) modularity optimization (FindNeighbors and FindClusters) at a resolution of 0.7 for LECs, 0.5 for B cells, and 0.1 for T cells. Gene expression was then normalized using *LogNormalize* (scale factor = 10,000) and log transformed. LEC identity was confirmed based on the expression of Prox1, and LEC subsets were further classified based on the expression of Ackr4, Foxc2, Lyve-1, Ptx3, Marco, Cd274, Madcam1, Ccl20^19^. Identity of GC B cells, plasma cells, and T_FH_ cells was confirmed based on the expression of the surface used for their initial sorting. Gene signature scores were calculated using *AddModuleScore*. Cell cycle phase assignment (G0– G1 vs. S–G2–M) was performed using the CellCycleScoring function and the cc.genes.updated.2019 dataset provided in Seurat. For replicate-level analyses of lymphocyte datasets, principal component analysis (PCA) was performed using SingleCellExperiment (v1.18.1) after applying a variance-stabilizing transformation with DESeq2 (v1.46.0). Batch effects were corrected using the removeBatchEffect function from limma (v3.62.2), and data visualization was performed with ggplot2.

### Gene set and pathway enrichment analysis

Gene set enrichment analysis (GSEA) was performed using fgsea (v1.32.4) with gene sets derived using escape v 2.5.5 from the “C2:CP:REACTOME” category for *Mus musculus*. Gene sets were generated from significantly differentially expressed genes (DEG) identified by comparing the same cell type across different genetic mouse lines with minimum size of 15. DEGs were ranked by average log2 fold change to generate ranked gene lists, which were subsequently sorted in descending order. Enrichment results were visualized using the enrichplot (v 1.26.6) and ggplot2(v 3.5.2).

### Immunoglobulin sequencing

Single GC B cells were index-sorted into 96-well plates containing 5 µl of TCL buffer (Qiagen) supplemented with 1% β-mercaptoethanol^16,37^. RNA was reverse-transcribed using oligo-dT as in ^16,19^. PCR primers were used as described previously^24^. Pooled PCR products were then purified using SPRI beads (0.7x volume ratio), gel purified and sequenced with a 600-cycle Reagent kit v3 on the Illumina Miseq platform.

### Immunoglobulin sequence analysis

Single-cell *Igh* analyses were carried out as described^24^. Raw, paired-end sequences were merged at the overlapping regions using PANDASeq (v 2.11) for full amplicon reconstruction, then processed with the FASTX toolkit. Only sequences with high counts for each single cell were analyzed. For V-(D)-J gene rearrangement annotation, *Ig* sequences obtained were submitted to HighV-QUEST (v. 1.9.5) and Vbase2 databases, choosing the assignment yielding the lowest number of somatic mutations in case of discrepancy. Sequences were then processed in the R environment to group cells with common V_H_/J_H_ genes and same CDR_H_3 length into clonal lineages when CDR_H_3 nucleotide identity was ≥ 75%. Thereafter, downstream mutational and clonality analyses were performed in Microsoft Excel.

### *In silico* germinal center modeling

We adapted a previously published simulator of the germinal center reaction to test the implications of follicle enlargement on the selection process. A comprehensive description of the simulator is provided in^25^ (MiXed-model). Modifications made to the original model are described below.

#### Space representation

The GC reaction space is represented as a three-dimensional discretized rectangular lattice with lattice constant of 𝛥𝑥 = 5𝜇m. Within the lattice, a spherical volume of radius 𝑟 defines the effective region in which cells are distributed and move. The sphere is conceptually divided into two zones, representing the DZ and LZ. Each lattice node can be occupied by a single cell only.

#### Space shape for antibodies

Antibodies are represented on a four (𝑑 = 4) dimensional shape space^77^. The shape space is restricted to a size of 10 positions per dimension, thus, only considering antibodies with a minimum affinity to the antigen. The optimal clone 𝜑* is positioned in the center of the shape space. A position on the shape space 𝜑 is attributed to each B cell. The 1-norm distance to the optimal clone, 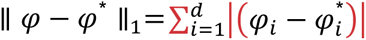, corresponds to the minimum number of mutations required to reach the optimal clone and is used as a measure for the antigen binding probability. The binding probability is calculated from the Gaussian distribution with width 𝛤 = 2.8 ^78^:

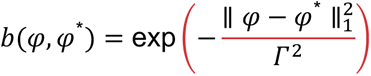

#### Founder cells

The model is initialized with a fixed number of T_FH_ cells, FDCs, and stromal cells. B cells enter the GC continuously at a defined rate per hour. For each simulated GC radius 𝑟, the total numbers of founder cells and the B cell influx rate are scaled by a volumetric factor (𝑟⁄𝑟_#$%_)^&^, where 𝑟_#$%_ = 160𝜇𝑚 is the reference radius corresponding to the wildtype condition. Approximate scaled values for each condition are as follows: for r160, T_FH_ cells, 140; FDCs, 200; stromal cells, 300; rate of B cel influx/ hour, 2.0; for r208, T_FH_ cells, 308; FDCs, 440; stromal cells, 661; rate of B cel influx/ hour, 4.4; for r240, T_FH_ cells, 473; FDCs, 676; stromal cells, 1013; rate of B cel influx/ hour, 6.76; for r320, T_FH_ cells, 1120; FDCs, 1600; stromal cells, 2400; rate of B cell influx/ hour, 16.0. T_FH_ cells are randomly distributed on the lattice and occupy a single node each. Stromal cells are restricted to the DZ, while FDCs are restricted to the LZ. FDCs are represented by their soma occupying one node and six dendrites that extend 40𝜇m from the soma. Dendrites are treated as transparent for B-cell or T_FH_-cell migration, while remaining accessible for B-cell interactions. Newly arriving B cells are randomly placed on unoccupied lattice nodes and assigned a shape space position, randomly selected from a predefined set of 100 positions.

#### Chemokine distribution

Two chemokines, CXCL12 and CXCL13, are included in the model. CXCL12 is produced by stromal cells in the DZ at a rate of 400nMol/h per stromal cell, and CXCL13 by FDCs in the LZ at a rate of 10nMol/h per FDC. As FDCs and stromal cells are immobile and constant in number within each simulation, the corresponding chemokine distributions were pre-calculated once for each considered GC radius 𝑟, and the resulting steady-state distributions were used throughout the simulation.

#### Implementation and data analysis

The simulation code was implemented in C++11 (v8.3). Simulation outputs were collected and processed in R (v3.5.2) for analysis and figure generation. The full code, parameter files and documentation are available on Zenodo^69^.

### Statistical analysis

Statistical analysis was performed using GraphPad Prism 10 software. Comparisons between two groups were conducted using unpaired Student’s t tests (parametric). The proper sample size was determined on an *ad hoc* basis drawing on prior experience. Gene set and significant pathways were identified by adjusting p-values to account for multiple testing using the Benjamini-Hochberg (BH) method. Statistical analysis of sequencing data was performed with pairwise Wilcoxon rank test and Bonferroni correction in R. P values are reported as exact numeric values.

## References

1. Liu, L., Zhong, Q., Tian, T., Dubin, K., Athale, S.K., and Kupper, T.S. (2010). Epidermal injury and infection during poxvirus immunization is crucial for the generation of highly protective T cell–mediated immunity. Nat Med 16, 224–227. 10.1038/nm.2078.

2. Hammarlund, E., Lewis, M.W., Hansen, S.G., Strelow, L.I., Nelson, J.A., Sexton, G.J., Hanifin, J.M., and Slifka, M.K. (2003). Duration of antiviral immunity after smallpox vaccination. Nat. Med. 9, 1131– 1137. 10.1038/nm917.

3. Lee, J.H., Sutton, H.J., Cottrell, C.A., Phung, I., Ozorowski, G., Sewall, L.M., Nedellec, R., Nakao, C., Silva, M., Richey, S.T., et al. (2022). Long-primed germinal centres with enduring affinity maturation and clonal migration. Nature 609, 998–1004. 10.1038/s41586-022-05216-9.

4. Hou, X., Zaks, T., Langer, R., and Dong, Y. (2021). Lipid nanoparticles for mRNA delivery. Nat Rev Mater 6, 1078–1094. 10.1038/s41578-021-00358-0.

5. Stritt, S., Koltowska, K., and Mäkinen, T. (2021). Homeostatic maintenance of the lymphatic vasculature. Trends Mol Med. 10.1016/j.molmed.2021.07.003.

6. Baluk, P., Fuxe, J., Hashizume, H., Romano, T., Lashnits, E., Butz, S., Vestweber, D., Corada, M., Molendini, C., Dejana, E., et al. (2007). Functionally specialized junctions between endothelial cells of lymphatic vessels. J. Exp. Med. 204, 2349–2362. 10.1084/jem.20062596.

7. Jalkanen, S., and Salmi, M. (2020). Lymphatic endothelial cells of the lymph node. Nat Rev Immunol, 1–13. 10.1038/s41577-020-0281-x.

8. Churchill, M.J., du Bois, H., Heim, T.A., Mudianto, T., Steele, M.M., Nolz, J.C., and Lund, A.W. (2022). Infection-induced lymphatic zippering restricts fluid transport and viral dissemination from skin. Journal of Experimental Medicine 219, e20211830. 10.1084/jem.20211830.

9. Loo, C.P., Nelson, N.A., Lane, R.S., Booth, J.L., Loprinzi Hardin, S.C., Thomas, A., Slifka, M.K., Nolz, J.C., and Lund, A.W. (2017). Lymphatic Vessels Balance Viral Dissemination and Immune Activation following Cutaneous Viral Infection. Cell Reports 20, 3176–3187. 10.1016/j.celrep.2017.09.006.

10. Victora, G.D., and Nussenzweig, M.C. (2022). Germinal Centers. Annu. Rev. Immunol. 40, 413–442. 10.1146/annurev-immunol-120419-022408.

11. Shih, T.-A.Y., Meffre, E., Roederer, M., and Nussenzweig, M.C. (2002). Role of BCR affinity in T cell dependent antibody responses in vivo. Nat Immunol 3, 570–575. 10.1038/ni803.

12. Bannard, O., and Cyster, J.G. (2017). Germinal centers: programmed for affinity maturation and antibody diversification. Curr Opin Immunol 45, 21–30. 10.1016/j.coi.2016.12.004.

13. Victora, G.D., Schwickert, T.A., Fooksman, D.R., Kamphorst, A.O., Meyer-Hermann, M., Dustin, M.L., and Nussenzweig, M.C. (2010). Germinal center dynamics revealed by multiphoton microscopy with a photoactivatable fluorescent reporter. Cell 143, 592–605. 10.1016/j.cell.2010.10.032.

14. Meyer-Hermann, M., Mohr, E., Pelletier, N., Zhang, Y., Victora, G.D., and Toellner, K.-M. (2012). A theory of germinal center B cell selection, division, and exit. Cell Rep 2, 162–174. 10.1016/j.celrep.2012.05.010.

15. Harrell, M.I., Iritani, B.M., and Ruddell, A. (2007). Tumor-induced sentinel lymph node lymphangiogenesis and increased lymph flow precede melanoma metastasis. Am. J. Pathol. 170, 774– 786. 10.2353/ajpath.2007.060761.

16. Angeli, V., Ginhoux, F., Llodrà, J., Quemeneur, L., Frenette, P.S., Skobe, M., Jessberger, R., Merad, M., and Randolph, G.J. (2006). B Cell-Driven Lymphangiogenesis in Inflamed Lymph Nodes Enhances Dendritic Cell Mobilization. Immunity 24, 203–215. 10.1016/j.immuni.2006.01.003.

17. Khan, T.N., Mooster, J.L., Kilgore, A.M., Osborn, J.F., and Nolz, J.C. (2016). Local antigen in nonlymphoid tissue promotes resident memory CD8+ T cell formation during viral infection. Journal of Experimental Medicine 213, 951–966. 10.1084/jem.20151855.

18. Gregory, J.L., Walter, A., Alexandre, Y.O., Hor, J.L., Liu, R., Ma, J.Z., Devi, S., Tokuda, N., Owada, Y., Mackay, L.K., et al. (2017). Infection Programs Sustained Lymphoid Stromal Cell Responses and Shapes Lymph Node Remodeling upon Secondary Challenge. Cell Reports 18, 406–418. 10.1016/j.celrep.2016.12.038.

19. Xiang, M., Grosso, R.A., Takeda, A., Pan, J., Bekkhus, T., Brulois, K., Dermadi, D., Nordling, S., Vanlandewijck, M., Jalkanen, S., et al. (2020). A Single-Cell Transcriptional Roadmap of the Mouse and Human Lymph Node Lymphatic Vasculature. Frontiers Cardiovasc Medicine 7, 52. 10.3389/fcvm.2020.00052.

20. Attaf, N., Baaklini, S., Binet, L., and Milpied, P. (2021). Heterogeneity of germinal center B cells: New insights from single-cell studies. Eur J Immunol 51, 2555–2567. 10.1002/eji.202149235.

21. Laidlaw, B.J., and Cyster, J.G. (2021). Transcriptional regulation of memory B cell differentiation. Nat Rev Immunol 21, 209–220. 10.1038/s41577-020-00446-2.

22. Kagan Ben Tikva, S., Gurwitz, N., Sivan, E., Hirsch, D., Hezroni-Barvyi, H., Biram, A., Moss, L., Wigoda, N., Egozi, A., Monziani, A., et al. (2024). T cell help induces Myc transcriptional bursts in germinal center B cells during positive selection. Sci Immunol 9, eadj7124. 10.1126/sciimmunol.adj7124.

23. Tas, J.M.J., Mesin, L., Pasqual, G., Targ, S., Jacobsen, J.T., Mano, Y.M., Chen, C.S., Weill, J.-C., Reynaud, C.-A., Browne, E.P., et al. (2016). Visualizing antibody affinity maturation in germinal centers. Science 351, 1048–1054. 10.1126/science.aad3439.

24. Nowosad, C.R., Mesin, L., Castro, T.B.R., Wichmann, C., Donaldson, G.P., Araki, T., Schiepers, A., Lockhart, A.A.K., Bilate, A.M., Mucida, D., et al. (2020). Tunable dynamics of B cell selection in gut germinal centres. Nature 588, 321–326. 10.1038/s41586-020-2865-9.

25. Meyer-Hermann, M. (2021). A molecular theory of germinal center B cell selection and division. Cell Reports 36, 109552. 10.1016/j.celrep.2021.109552.

26. Calado, D.P., Sasaki, Y., Godinho, S.A., Pellerin, A., Köchert, K., Sleckman, B.P., de Alborán, I.M., Janz, M., Rodig, S., and Rajewsky, K. (2012). The cell-cycle regulator c-Myc is essential for the formation and maintenance of germinal centers. Nat Immunol 13, 1092–1100. 10.1038/ni.2418.

27. Dominguez-Sola, D., Victora, G.D., Ying, C.Y., Phan, R.T., Saito, M., Nussenzweig, M.C., and Dalla-Favera, R. (2012). The proto-oncogene MYC is required for selection in the germinal center and cyclic reentry. Nat Immunol 13, 1083–1091. 10.1038/ni.2428.

28. Shulman, Z., Gitlin, A.D., Weinstein, J.S., Lainez, B., Esplugues, E., Flavell, R.A., Craft, J.E., and Nussenzweig, M.C. (2014). Dynamic signaling by T follicular helper cells during germinal center B cell selection. Science 345, 1058–1062. 10.1126/science.1257861.

29. Crotty, S. (2011). Follicular helper CD4 T cells (TFH). Annu Rev Immunol 29, 621–663. 10.1146/annurev-immunol-031210-101400.

30. Schwickert, T.A., Victora, G.D., Fooksman, D.R., Kamphorst, A.O., Mugnier, M.R., Gitlin, A.D., Dustin, M.L., and Nussenzweig, M.C. (2011). A dynamic T cell-limited checkpoint regulates affinity-dependent B cell entry into the germinal center. J Exp Med 208, 1243–1252. 10.1084/jem.20102477.

31. Weinstein, J.S., Herman, E.I., Lainez, B., Licona-Limón, P., Esplugues, E., Flavell, R., and Craft, J. (2016). TFH cells progressively differentiate to regulate the germinal center response. Nat Immunol 17, 1197–1205. 10.1038/ni.3554.

32. Gonzalez, D.G., Cote, C.M., Patel, J.R., Smith, C.B., Zhang, Y., Nickerson, K.M., Zhang, T., Kerfoot, S.M., and Haberman, A.M. (2018). Nonredundant Roles of IL-21 and IL-4 in the Phased Initiation of Germinal Center B Cells and Subsequent Self-Renewal Transitions. J Immunol 201, 3569–3579. 10.4049/jimmunol.1500497.

33. Dalit, L., Tan, C.W., Sheikh, A.A., Munnings, R., Howson, L.J., Alvarado, C., Hussain, T., Zaini, A., Cooper, L., Kirn, A., et al. (2025). Divergent cytokine and transcriptional signatures control functional T follicular helper cell heterogeneity. Nat Immunol 26, 1821–1835. 10.1038/s41590-025-02258-9.

34. Podestà, M.A., Cavazzoni, C.B., Hanson, B.L., Bechu, E.D., Ralli, G., Clement, R.L., Zhang, H., Chandrakar, P., Lee, J.-M., Reyes-Robles, T., et al. (2023). Stepwise differentiation of follicular helper T cells reveals distinct developmental and functional states. Nat Commun 14, 7712. 10.1038/s41467-023-43427-4.

35. Merkenschlager, J., Finkin, S., Ramos, V., Kraft, J., Cipolla, M., Nowosad, C.R., Hartweger, H., Zhang, W., Olinares, P.D.B., Gazumyan, A., et al. (2021). Dynamic regulation of TFH selection during the germinal centre reaction. Nature 591, 458–463. 10.1038/s41586-021-03187-x.

36. Petersone, L., and Walker, L.S.K. (2024). T-cell help in the germinal center: homing in on the role of IL-21. Int Immunol 36, 89–98. 10.1093/intimm/dxad056.

37. Pae, J., Ersching, J., Castro, T.B.R., Schips, M., Mesin, L., Allon, S.J., Ordovas-Montanes, J., Mlynarczyk, C., Melnick, A., Efeyan, A., et al. (2021). Cyclin D3 drives inertial cell cycling in dark zone germinal center B cells. J Exp Med 218, e20201699. 10.1084/jem.20201699.

38. Shulman, Z., Gitlin, A.D., Targ, S., Jankovic, M., Pasqual, G., Nussenzweig, M.C., and Victora, G.D. (2013). T Follicular Helper Cell Dynamics in Germinal Centers. Science 341, 673–677. 10.1126/science.1241680.

39. Lucas, E.D., Finlon, J.M., Burchill, M.A., McCarthy, M.K., Morrison, T.E., Colpitts, T.M., and Tamburini, B.A.J. (2018). Type 1 IFN and PD-L1 Coordinate Lymphatic Endothelial Cell Expansion and Contraction during an Inflammatory Immune Response. J.I. 201, 1735–1747. 10.4049/jimmunol.1800271.

40. Kataru, R.P., Kim, H., Jang, C., Choi, D.K., Koh, B.I., Kim, M., Gollamudi, S., Kim, Y.-K., Lee, S.-H., and Koh, G.Y. (2011). T lymphocytes negatively regulate lymph node lymphatic vessel formation. Immunity 34, 96–107. 10.1016/j.immuni.2010.12.016.

41. Karakousi, T., Cristaldi, V., Lopes de Oliveira, M.L., Medeiros Geraldo, L.H., Gonzalez-Robles, T., da Silva, G., Breazeale, A.P., Encarnacion Rosado, J., Pozniak, J., Kimmelman, A., et al. (2024). IFNγ-dependent metabolic reprogramming restrains an immature, pro-metastatic lymphatic state in melanoma. bioRxiv, 2024.12.02.626426. 10.1101/2024.12.02.626426.

42. Petkova, M., Kraft, M., Stritt, S., Martinez-Corral, I., Ortsäter, H., Vanlandewijck, M., Jakic, B., Baselga, E., Castillo, S.D., Graupera, M., et al. (2023). Immune-interacting lymphatic endothelial subtype at capillary terminals drives lymphatic malformation. Journal of Experimental Medicine 220, e20220741. 10.1084/jem.20220741.

43. Junt, T., Moseman, E.A., Iannacone, M., Massberg, S., Lang, P.A., Boes, M., Fink, K., Henrickson, S.E., Shayakhmetov, D.M., Paolo, N.C.D., et al. (2007). Subcapsular sinus macrophages in lymph nodes clear lymph-borne viruses and present them to antiviral B cells. Nature 450, 110–114. 10.1038/nature06287.

44. Zhang, F., Zarkada, G., Han, J., Li, J., Dubrac, A., Ola, R., Genet, G., Boyé, K., Michon, P., Künzel, S.E., et al. (2018). Lacteal junction zippering protects against diet-induced obesity. Science 361, 599– 603. 10.1126/science.aap9331.

45. Kähäri, L., Fair-Mäkelä, R., Auvinen, K., Rantakari, P., Jalkanen, S., Ivaska, J., and Salmi, M. (2019). Transcytosis route mediates rapid delivery of intact antibodies to draining lymph nodes. J Clin Invest 129, 3086–3102. 10.1172/JCI125740.

46. Triacca, V., Güç, E., Kilarski, W.W., Pisano, M., and Swartz, M.A. (2017). Transcellular Pathways in Lymphatic Endothelial Cells Regulate Changes in Solute Transport by Fluid Stress. Circulation Research 120, 1440–1452. 10.1161/CIRCRESAHA.116.309828.

47. Kwon, D.S., Gregorio, G., Bitton, N., Hendrickson, W.A., and Littman, D.R. (2002). DC-SIGN-mediated internalization of HIV is required for trans-enhancement of T cell infection. Immunity 16, 135–144. 10.1016/s1074-7613(02)00259-5.

48. Qi, H., Egen, J.G., Huang, A.Y.C., and Germain, R.N. (2006). Extrafollicular activation of lymph node B cells by antigen-bearing dendritic cells. Science 312, 1672–1676. 10.1126/science.1125703.

49. Cyster, J.G. (2010). B cell follicles and antigen encounters of the third kind. Nat Immunol 11, 989–996. 10.1038/ni.1946.

50. Harwood, N.E., and Batista, F.D. (2010). Early Events in B Cell Activation. Annu. Rev. Immunol. 28, 185–210. 10.1146/annurev-immunol-030409-101216.

51. Phan, T.G., Grigorova, I., Okada, T., and Cyster, J.G. (2007). Subcapsular encounter and complement-dependent transport of immune complexes by lymph node B cells. Nat Immunol 8, 992–1000. 10.1038/ni1494.

52. Nowosad, C.R., Spillane, K.M., and Tolar, P. (2016). Germinal center B cells recognize antigen through a specialized immune synapse architecture. Nat Immunol 17, 870–877. 10.1038/ni.3458.

53. Khalil, A.M., Cambier, J.C., and Shlomchik, M.J. (2012). B cell receptor signal transduction in the GC is short-circuited by high phosphatase activity. Science 336, 1178–1181. 10.1126/science.1213368.

54. Spillane, K.M., and Tolar, P. (2017). B cell antigen extraction is regulated by physical properties of antigen-presenting cells. Journal of Cell Biology 216, 217–230. 10.1083/jcb.201607064.

55. Tam, H.H., Melo, M.B., Kang, M., Pelet, J.M., Ruda, V.M., Foley, M.H., Hu, J.K., Kumari, S., Crampton, J., Baldeon, A.D., et al. (2016). Sustained antigen availability during germinal center initiation enhances antibody responses to vaccination. Proc. Natl. Acad. Sci. U.S.A. 113. 10.1073/pnas.1606050113.

56. Cirelli, K.M., Carnathan, D.G., Nogal, B., Martin, J.T., Rodriguez, O.L., Upadhyay, A.A., Enemuo, C.A., Gebru, E.H., Choe, Y., Viviano, F., et al. (2019). Slow Delivery Immunization Enhances HIV Neutralizing Antibody and Germinal Center Responses via Modulation of Immunodominance. Cell 177, 1153–1171.e28. 10.1016/j.cell.2019.04.012.

57. Pauthner, M., Havenar-Daughton, C., Sok, D., Nkolola, J.P., Bastidas, R., Boopathy, A.V., Carnathan, D.G., Chandrashekar, A., Cirelli, K.M., Cottrell, C.A., et al. (2017). Elicitation of Robust Tier 2 Neutralizing Antibody Responses in Nonhuman Primates by HIV Envelope Trimer Immunization Using Optimized Approaches. Immunity 46, 1073–1088.e6. 10.1016/j.immuni.2017.05.007.

58. Pereira, J.P., Kelly, L.M., and Cyster, J.G. (2010). Finding the right niche: B-cell migration in the early phases of T-dependent antibody responses. Int Immunol 22, 413–419. 10.1093/intimm/dxq047.

59. Lu, E., and Cyster, J.G. (2019). G-protein coupled receptors and ligands that organize humoral immune responses. Immunol Rev 289, 158–172. 10.1111/imr.12743.

60. Chorny, A., Casas-Recasens, S., Sintes, J., Shan, M., Polentarutti, N., García-Escudero, R., Walland, A.C., Yeiser, J.R., Cassis, L., Carrillo, J., et al. (2016). The soluble pattern recognition receptor PTX3 links humoral innate and adaptive immune responses by helping marginal zone B cells. J Exp Med 213, 2167–2185. 10.1084/jem.20150282.

61. Turati, M., Giacomini, A., Rezzola, S., Maccarinelli, F., Gazzaroli, G., Valentino, S., Bottazzi, B., Presta, M., and Ronca, R. (2022). The natural FGF-trap long pentraxin 3 inhibits lymphangiogenesis and lymphatic dissemination. Exp Hematol Oncol 11, 84. 10.1186/s40164-022-00330-w.

62. Rovere, P., Peri, G., Fazzini, F., Bottazzi, B., Doni, A., Bondanza, A., Zimmermann, V.S., Garlanda, C., Fascio, U., Sabbadini, M.G., et al. (2000). The long pentraxin PTX3 binds to apoptotic cells and regulates their clearance by antigen-presenting dendritic cells. Blood 96, 4300–4306. 10.1182/blood.V96.13.4300.

63. Ruddell, A., Harrell, M.I., Furuya, M., Kirschbaum, S.B., and Iritani, B.M. (2011). B Lymphocytes Promote Lymphogenous Metastasis of Lymphoma and Melanoma. Neoplasia 13, 748–757. 10.1593/neo.11756.

64. Clatworthy, M.R., Harford, S.K., Mathews, R.J., and Smith, K.G.C. (2014). FcγRIIb inhibits immune complex-induced VEGF-A production and intranodal lymphangiogenesis. Proc National Acad Sci 111, 17971–17976. 10.1073/pnas.1413915111.

65. de Carvalho, R.V.H., Ersching, J., Barbulescu, A., Hobbs, A., Castro, T.B.R., Mesin, L., Jacobsen, J.T., Phillips, B.K., Hoffmann, H.-H., Parsa, R., et al. (2023). Clonal replacement sustains long-lived germinal centers primed by respiratory viruses. Cell 186, 131–146.e13. 10.1016/j.cell.2022.11.031.

66. Degn, S.E., van der Poel, C.E., Firl, D.J., Ayoglu, B., Al Qureshah, F.A., Bajic, G., Mesin, L., Reynaud, C.-A., Weill, J.-C., Utz, P.J., et al. (2017). Clonal Evolution of Autoreactive Germinal Centers. Cell 170, 913–926.e19. 10.1016/j.cell.2017.07.026.

67. Vinuesa, C.G., Tangye, S.G., Moser, B., and Mackay, C.R. (2005). Follicular B helper T cells in antibody responses and autoimmunity. Nat Rev Immunol 5, 853–865. 10.1038/nri1714.

68. Vinuesa, C.G., Sanz, I., and Cook, M.C. (2009). Dysregulation of germinal centres in autoimmune disease. Nat Rev Immunol 9, 845–857. 10.1038/nri2637.

69. Schips, M., Schwan, N., and Meyer-Hermann, M. (2025). GC SIMULATION: Lymphatic constraint of germinal centers optimizes protective antibody responses. (Zenodo). 10.5281/ZENODO.17494281 10.5281/ZENODO.17494281.

70. Bazigou, E., Lyons, O.T.A., Smith, A., Venn, G.E., Cope, C., Brown, N.A., and Makinen, T. (2011). Genes regulating lymphangiogenesis control venous valve formation and maintenance in mice. J. Clin. Invest. 121, 2984–2992. 10.1172/JCI58050.

71. Dogan, I., Bertocci, B., Vilmont, V., Delbos, F., Mégret, J., Storck, S., Reynaud, C.-A., and Weill, J.-C. (2009). Multiple layers of B cell memory with different effector functions. Nat Immunol 10, 1292–1299. 10.1038/ni.1814.

72. Snippert, H.J., van der Flier, L.G., Sato, T., van Es, J.H., van den Born, M., Kroon-Veenboer, C., Barker, N., Klein, A.M., van Rheenen, J., Simons, B.D., et al. (2010). Intestinal crypt homeostasis results from neutral competition between symmetrically dividing Lgr5 stem cells. Cell 143, 134–144. 10.1016/j.cell.2010.09.016.

73. Wyatt, L.S., Earl, P.L., and Moss, B. (2017). Generation of Recombinant Vaccinia Viruses. Curr Protoc Protein Sci 89, 5.13.1-5.13.18. 10.1002/cpps.33.

74. Oldstone, M.B., Tishon, A., Eddleston, M., de la Torre, J.C., McKee, T., and Whitton, J.L. (1993). Vaccination to prevent persistent viral infection. J Virol 67, 4372–4378. 10.1128/JVI.67.7.4372-4378.1993.

75. Shen, H., Miller, J.F., Fan, X., Kolwyck, D., Ahmed, R., and Harty, J.T. (1998). Compartmentalization of Bacterial Antigens: Differential Effects on Priming of CD8 T Cells and Protective Immunity. Cell 92, 535–545. 10.1016/S0092-8674(00)80946-0.

76. Korsunsky, I., Millard, N., Fan, J., Slowikowski, K., Zhang, F., Wei, K., Baglaenko, Y., Brenner, M., Loh, P., and Raychaudhuri, S. (2019). Fast, sensitive and accurate integration of single-cell data with Harmony. Nat Methods 16, 1289–1296. 10.1038/s41592-019-0619-0.

77. Perelson, A.S., and Oster, G.F. (1979). Theoretical studies of clonal selection: minimal antibody repertoire size and reliability of self-non-self discrimination. J Theor Biol 81, 645–670. 10.1016/0022-5193(79)90275-3.

78. Meyer-Hermann, M., Deutsch, A., and Or-Guil, M. (2001). Recycling probability and dynamical properties of germinal center reactions. J Theor Biol 210, 265–285. 10.1006/jtbi.2001.2297.

