## Supplemental Figures for "Lymphatic constraint of germinal centers optimizes protective antibody responses"

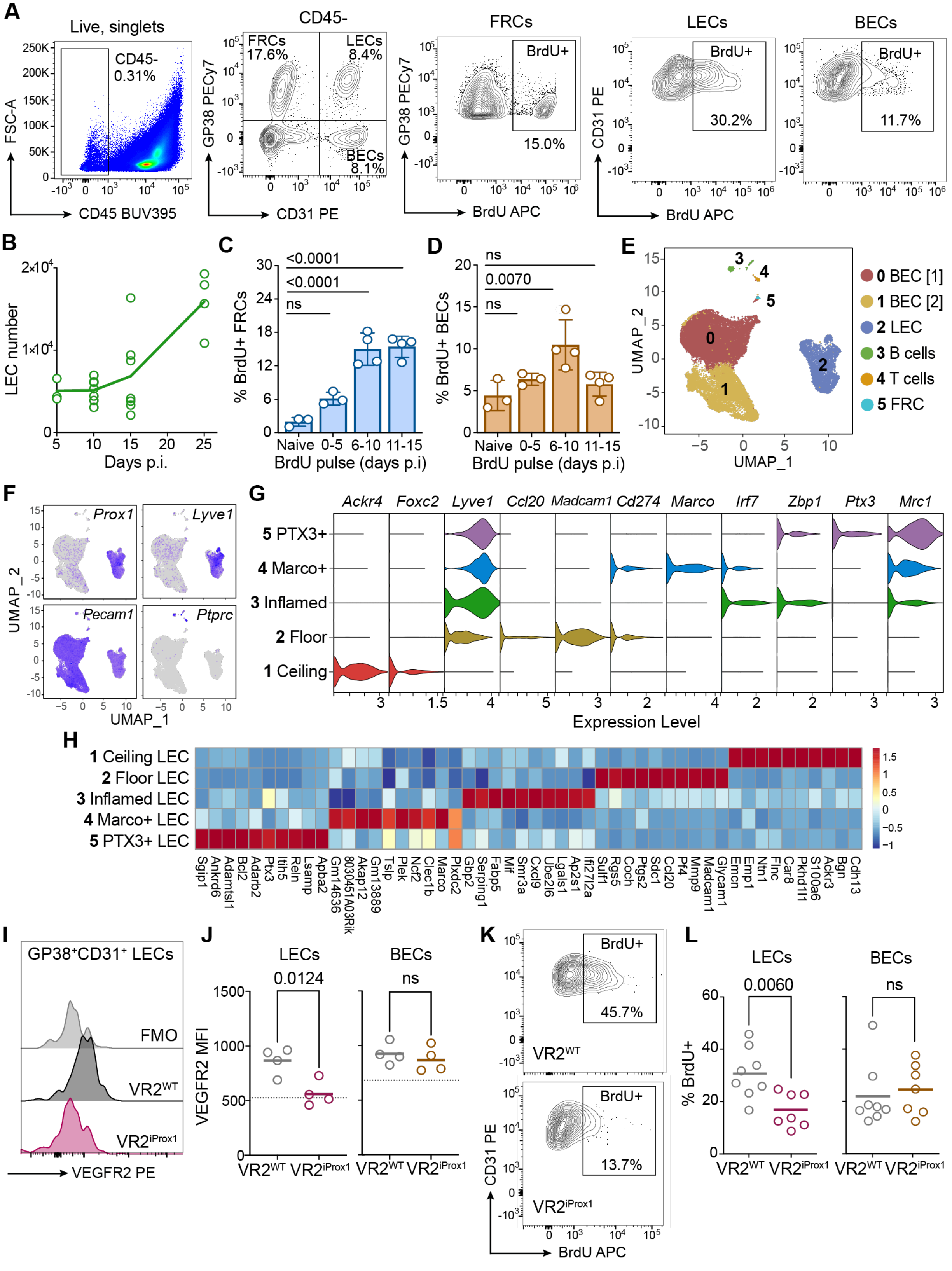


**Supplemental Figure 1. Stromal and lymphatic endothelial cell subset composition and proliferation dynamics in dLNs of WT and LEC VEGFR2-deficient mice following VACV scarification.**

(**A**) Flow cytometry gating strategy to identify lymphatic endothelial cells (LECs; $CD45^{-}CD31^{+}gp38^{+}$), blood endothelial cells (BECs; $CD45^{-}CD31^{+}gp38^{-}$), fibroblastic reticular cells (FRCs; $CD45^{-}CD31^{-}gp38^{+}$), and the proliferating ($BrdU^{+}$) subsets in the draining lymph nodes (dLN).

(**B**) Total number of LECs per dLN at indicated days after vaccinia virus (VACV) infection (by scarification), quantified by flow cytometry. Each point is one mouse.

(**C-D**) Quantification of percent BrdU$^+$ FRCs (C) and BECs (D) of parent population in the dLNs after indicated BrdU pulse intervals (5 days) following VACV infection. *p* values calculated using one way ANOVA.

**(E)** UMAP projection of scRNA-seq of pooled endothelial-enriched cells (CD45$^-$CD31$^+$) sorted from dLNs of C57BL/6 wildtype mice at days 0, 5, 7, 15, and 25 post VACV infection. Clusters are color-coded by subset identity: BEC[1] (0), BEC[2] (1), LEC (2), B cells (3), T cells (4), FRC (5).

**(F)** Expression of key genes expressed by clusters shown in (E): lymphatic endothelial cells (LECs, *Prox1*, *Lyve1*), endothelial cells (*Pecam1*), and leukocytes (*Ptprc*).

(**G**) Expression of cluster-defining genes, subset from (E) and re-clustered.

(**H**) Heatmap of top 10 marker genes by LEC cluster. Each column represents the relative z-score of expression of classifier genes, and each row describes a LEC subset (as labeled). Color scale indicates the $\log_2$ fold change of average expression.

(**I and J**) Representative histogram (I) and quantification (J) of mean fluorescence intensity (MFI) of VEGFR2 on LECs and BECs in dLNs of VEGFR2$^{WT}$ control and VEGFR2$^{iProx1}$ mice at day 5 post VACV infection. Fluorescence-minus-one (FMO) control is indicated by dotted lines. Each point is one mouse; lines indicate median; *p* values were calculated using two-tailed Student's *t* test. Data are representative of two independent experiments. ns= not significant.

(**K and L**) Representative flow cytometry plots (K) and quantification (L) of BrdU$^+$ LN LECs and BECs from dLNs of VEGFR2$^{WT}$ control and VEGFR2$^{iProx1}$ mice pulsed with BrdU at days 0-5 post VACV infection. Each point is one mouse; lines indicate median; *p* values were calculated using two-tailed Student's *t* test. Data are from two independent experiments (n=8 mice, VEGFR2$^{WT}$; n=7 mice, VEGR2$^{iProx1}$).

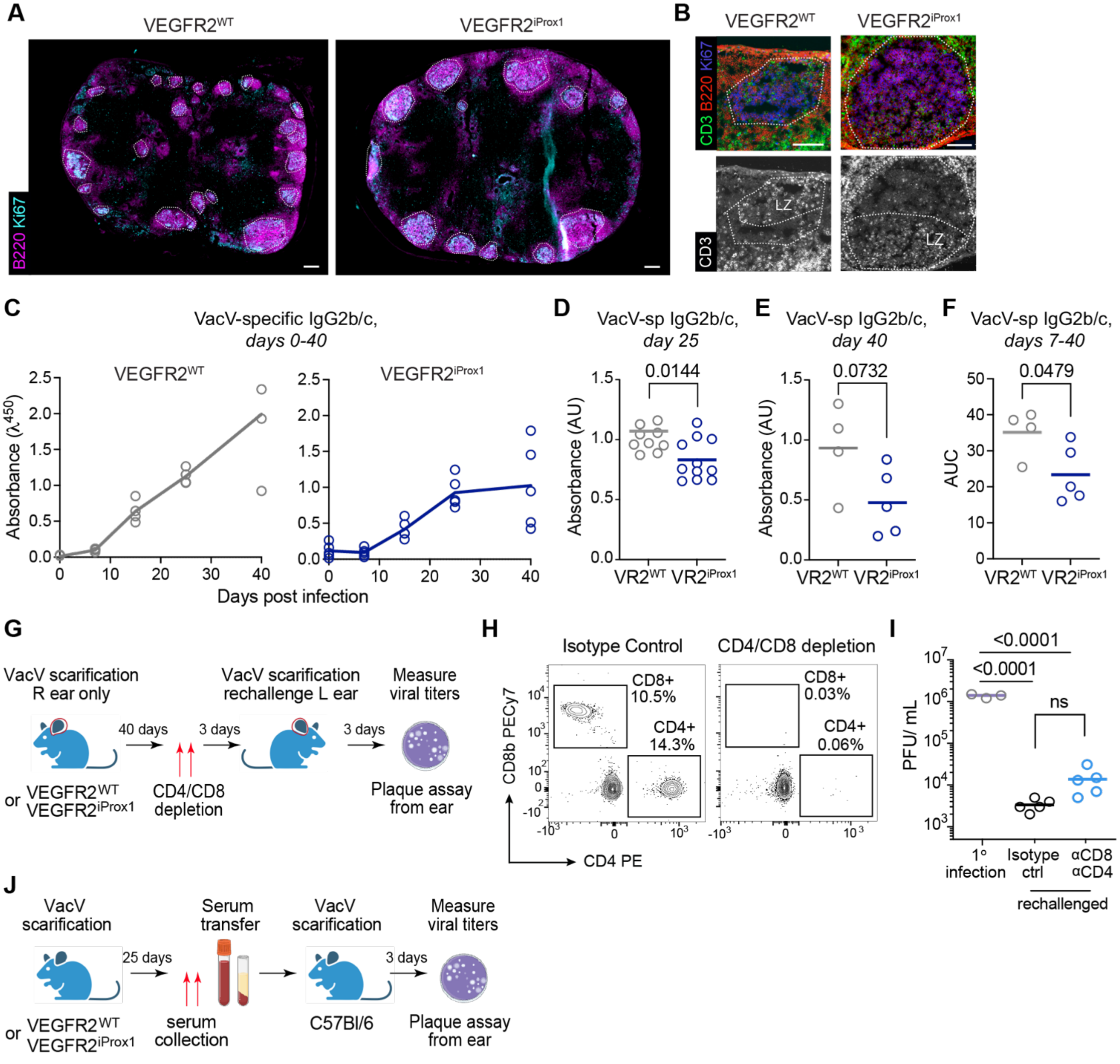


**Supplemental Figure 2. VACV-specific antibody responses in VEGFR2$^{WT}$ and VEGFR2$^{iProx1}$ mice.**

(**A**) Representative immunofluorescence images of dLN cross sections from VEGFR2$^{WT}$ (VR2$_{WT}$) and VEGFR2$^{iProx1}$ (VR2$^{iProx1}$) mice showing germinal centers (GC) at day 15 post vaccinia virus (VACV) infection (by scarification). Magenta, B220; cyan, Ki-67. Scale bar = 200 μm.

(**B**) Representative immunofluorescence images of GCs (top) in VEGFR2$^{WT}$ and VEGFR2$^{iProx1}$ mice at day 15 post-VACV infection. Red, B220; blue, Ki-67; green, CD3ε. Dotted lines outline GCs (Ki-67 area). Scale bars = 50 μm. Representative image (bottom) of CD3ε channel only, showing GC T cells (T$_{FH}$ cells) in the GC light zone (LZ). Dotted lines indicate GC and LZ boundaries.

(**C**) Longitudinal analysis of VACV-specific antibody responses. Serum from VEGFR2$^{WT}$ (left) and VEGR2$^{iProx1}$ (right) mice infected with VACV by scarification was collected at indicated time points, and VACV-specific IgG2b/c was quantified by ELISA (n=3 mice, VEGFR2$^{WT}$; n=5 mice, VEGR2$^{iProx1}$). Each point is one mouse; lines connect mean values across time points.

(**D and E**) VACV-specific serum IgG2b/c from VEGFR2$^{WT}$ and VEGFR2$^{iProx1}$ mice at day 25 (D) or day 40 (E) post VACV infection, assessed by ELISA. Each point is one mouse; lines indicate median. Data are from four independent experiments (n=13 mice, VEGFR2$^{WT}$; n=15 mice, VEGR2$^{iProx1}$) (D) and one experiment (n=4 mice, VEGFR2$^{WT}$; n=5 mice, VEGR2$^{iProx1}$) (E); *p* values were calculated using two-tailed Student's *t* test.

(**F**) Area under the curve quantification of VACV-specific IgG2b/c antibodies overtime of VEGFR2$^{WT}$ and VEGR2$^{iProx1}$ in (A). Each point is one mouse; lines indicate median values. Data are from one experiment (n=4 mice, VEGFR2$^{WT}$; n=5 mice, VEGFR2$^{iProx1}$); *p* values were calculated using two-tailed Student's *t* test.

(**G**) Schematic of the VACV re-challenge experiment.

(**H**) Representative flow cytometry plots of blood from mice treated with $\alpha$CD8 and $\alpha$CD4 depletion antibodies or isotype control.

(**I**) Viral titers (Plaque-forming unites, PFU) in ears of C57BL/6 wildtype mice were measured following primary VACV infection, VACV rechallenge with isotype control treatment, and VACV rechallenge after $\alpha$CD8 and $\alpha$CD4 depletion.

(**J**) Schematic of the VACV serum transfer experiment.

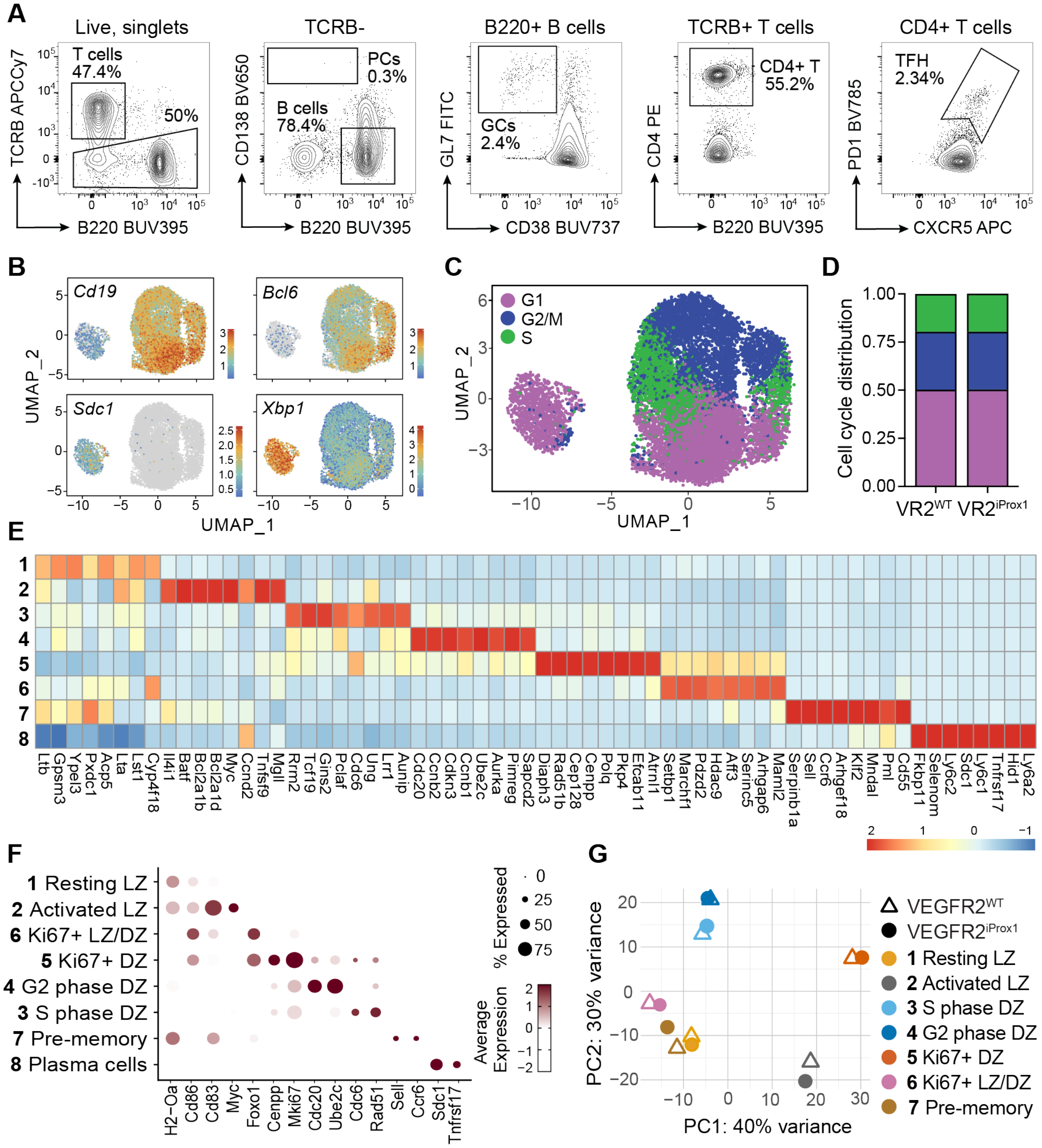


**Supplemental Figure 3 GC B cell subsets in VEGFR2$^{WT}$ and VEGFR2$^{iProx1}$ mice.**

(**A**) Flow cytometry gating strategy for FACS sorting of GC B cells (TCRβ$^{-}$B220$^{+}$GL7$^{+}$CD38$^{-}$), plasma cells (PCs; TCRβ$^{-}$B220$^{mid-lo}$CD138$^{+}$), and T$_{FH}$ cells (TCRβ$^{+}$B220$^{-}$CD4$^{+}$PD1$^{+}$CXCR5$^{+}$) from dLNs of VEGFR2$^{WT}$ (VR2$^{WT}$) and VEGFR2$^{iProx1}$ (VR2$^{iProx1}$) mice at day 25 post-VACV scarification used for downstream scRNA-seq analysis.

**(B)** Expression levels of key marker genes used to define B cell clusters; B cells (*Cd19*), GC B cells (*Bcl6*), PCs (*Xbp1*, *Sdc1*).

(**C**) UMAP projection of B cell scRNA-seq color coded based on the expression of genes associated with the S (green)- G2/M (blue)- G1 (purple) stages of the cell cycle.

(**D**) Proportional distribution of each cell cluster in (C) separated by genotype VEGFR2$^{WT}$ and VEGFR2$^{iProx1}$.

(**E**) Heatmap summarizing the average expression of B cell cluster defining genes (top 8). Each numbered row represents B cell subsets as defined in (F), and each column shows the relative z-scored gene expression. Color scale indicates the log$_2$ fold change of average expression.

(**F**) Expression of B cell cluster defining genes. Dot size denotes the proportion of cells expressing a given gene, color intensity denotes the relative average expression level of the corresponding gene, using the original log-transformed counts.

(**G**) PCA plot of GC B cell clusters. Cluster 8, plasma cells, was excluded to improve resolution amongst GC clusters. Color denotes subcluster, shape denotes genotype. Triangles indicate VEGFR2$^{WT}$ and circles indicate VEGFR2$^{iProx1}$.

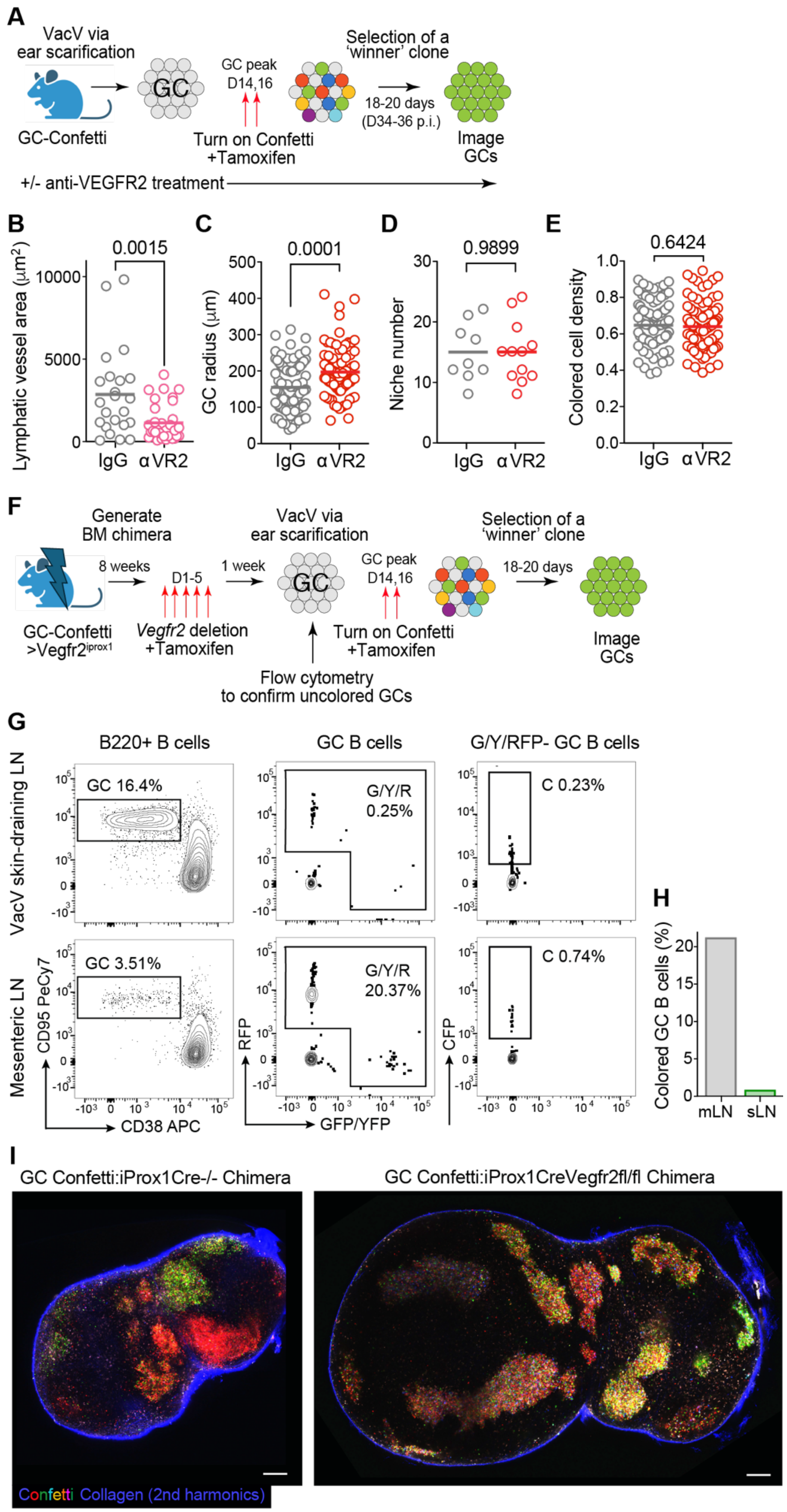

A
VacV via ear scarification
GC-Confetti
GC
GC peak D14,16
Turn on Confetti +Tamoxifen
Selection of a 'winner' clone
18-20 days (D34-36 p.i.)
Image GCs
+/- anti-VEGFR2 treatment
B
Lymphatic vessel area (μm²)
0.0015
IgG
αVR2
C
GC radius (μm)
0.0001
D
Niche number
0.9899
E
Colored cell density
0.6424
F
Generate BM chimera
8 weeks
D1-5
1 week
VacV via ear scarification
GC peak D14,16
Selection of a 'winner' clone
18-20 days
GC-Confetti >Vegfr2iprox1
Vegfr2 deletion +Tamoxifen
Flow cytometry to confirm uncolored GCs
Turn on Confetti +Tamoxifen
Image GCs
G
B220+ B cells
GC B cells
G/Y/RFP- GC B cells
VacV skin-draining LN
GC 16.4%
G/Y/R 0.25%
C 0.23%
Mesenteric LN
GC 3.51%
G/Y/R 20.37%
C 0.74%
CD95 PeCy7
CD38 APC
RFP
GFP/YFP
CFP
H
Colored GC B cells (%)
mLN
sLN
I
GC Confetti:iProx1Cre-/- Chimera
GC Confetti:iProx1CreVegfr2fl/fl Chimera
Confetti Collagen (2nd harmonics)

**Supplemental Figure 4. Experimental design and control validation for GC selection studies.**

(**A**) Schematic describing use of the Confetti allele tracking selection *in vivo* in vaccinia virus (VACV)-induced germinal centers (GCs).

(**B**) Quantification of perifollicular lymphatic vessel area (LYVE1) surrounding individual B follicles in the draining lymph nodes (dLNs) of C57BL/6 wildtype mice treated with IgG isotype control or anti-VEGFR2 antibodies ($\alpha$VR2) at day 5 post-VACV infection (by scarification). Each symbol represents a single B cell follicle. Data are from one experiment (n = 3 IgG, n = 4 $\alpha$VR2). Lines indicate the median; p values were calculated using a two-tailed Student's t test.

(**C-E**) Quantification of GC radius (C), GC niche number (D), and colored cell density (E) from AID-Confetti GCs in mice treated with IgG isotype control or $\alpha$VR2 imaged with multiphoton microscopy. Each point is a GC. Data are from 2 independent experiments (n = 7 mice, IgG, n = 6 mice, $\alpha$VR2). Lines indicate the median; p values were calculated using a two-tailed Student's t test.

(**F**) Schematic describing the deletion of VEGFR2 and the induction of the Confetti allele in bone marrow chimera experiments.

(**G**) Representative flow cytometry plots showing the frequency of confetti-labeled GC B cells ($CD19^{+}B220^{+}CD95^{+}CD38^{-}$RFP/GFP/YFP/$CFP^{+}$) in chimeric mice in either skin-draining (sLN) or mesenteric LNs (mLN) following the first dose of tamoxifen (time point as indicated by first set of red arrows in (F). Note that initial tamoxifen treatment targeting VEGFR2 induces Confetti labeling in constitutive mesenteric GCs, but spares VACV GCs induced after VEGFR2 deletion.

(**H**) Quantification of GC Confetti labeling shown in (G) in either 1 mLN or 2 sLNs.

(**I**) Representative multiphoton images of dLNs from control or VEGFR2 deficient chimeric mice as indicated in (F). Scale bars = 200 μm.

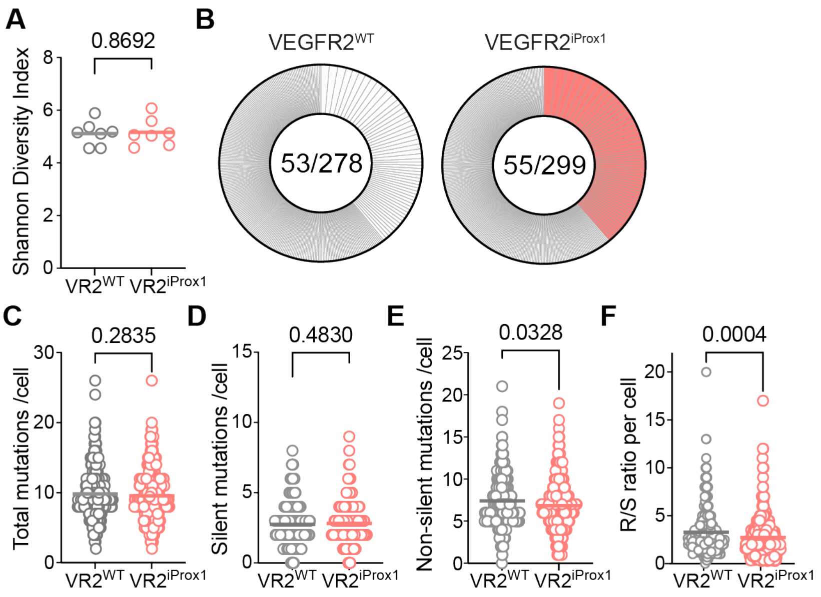


**Supplemental Figure 5. GC B cell clonal diversity and somatic hypermutation in VEGFR2$^{WT}$ and VEGFR2$^{iProx1}$ mice following VACV infection.**

(**A**) Shannon diversity index of B cell clones derived from single-cell V(D)J sequencing of the Ig heavy chain of individual sorted germinal center (GC) B cells (B220$^{+}$CD38$^{-}$ GL-7$^{+}$) from VEGFR2$^{WT}$ (VR2$^{WT}$) and VEGFR2$^{iProx1}$ (VR2$^{iProx1}$) mice at day 25 post-VACV infection (by scarification). Each point is one mouse (33-135 cells per mouse); lines indicate mean. Data are from two independent experiments (n=7 mice, VEGFR2$^{WT}$; n=7 mice, VEGR2$^{iProx1}$); p values were calculated using a two-tailed Student's t test.

(**B**) Analysis of GC IgH clonal expansions using data from (A). Colored/ white segments are expanded clones, where width of segment indicates degree of expansion. Narrow segments indicate singlets. Numbers in pie charts are number of clones out of the total number of cells sequenced.

(**C-F**) Mutational analysis of data from (A) showing total (C), silent (D), and non-silent (E), mutations per cell. (F) ratio of replacement (non-silent) to silent mutations per cell. Each point is an individual cell; lines indicate the mean. p values were calculated using a two-tailed Student's t test.

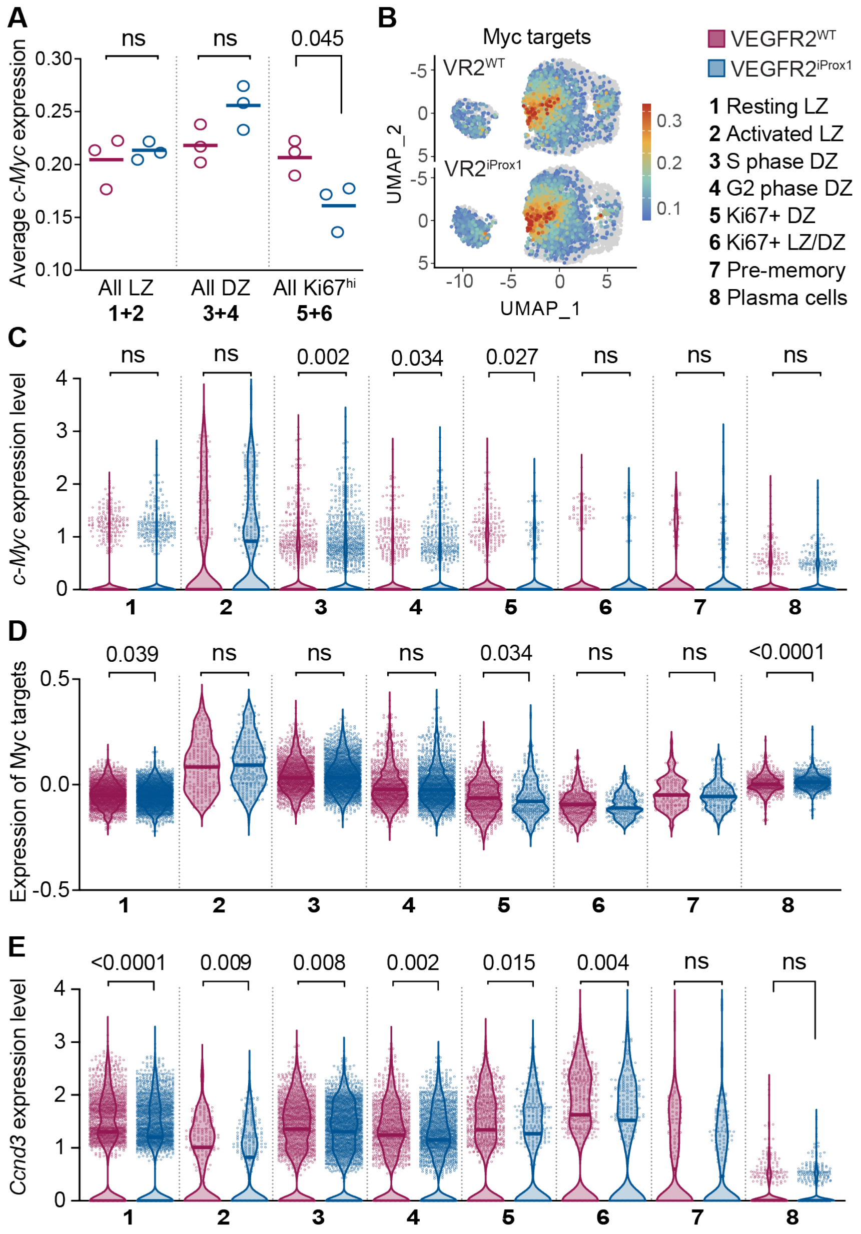


**Supplemental Figure 6. C-Myc, Myc target genes, and ccnd3 expression activity in GC B cell subsets of VEGFR2$^{WT}$ and VEGFR2$^{iProx1}$ mice.**

(**A**) Average *c-Myc* expression across pooled B cell subclusters (numbers correspond to annotations Figure 3C), comparing VEGFR2$^{WT}$ (purple, VR2$^{WT}$) and VEGFR2$^{iProx1}$ (blue, VR2$^{iProx1}$). Each point is one mouse; lines indicate median; p values were calculated using a two-tailed Student's t test.

(**B**) UMAP projection of B cell scRNA-seq data split by genotype (VEGFR2$^{WT}$ control and VEGFR2$^{iProx1}$) and colored by z-scored expression of Myc target genes.

(**C-E**) Relative expression of *c-Myc* (C), Myc target genes (D), and *Ccnd3* (E) in individual cells across B cell subsets, in VEGFR2$^{WT,}$ purple; or VEGFR2$^{iProx1,}$ blue. p values were calculated using Mann-Whitney test.

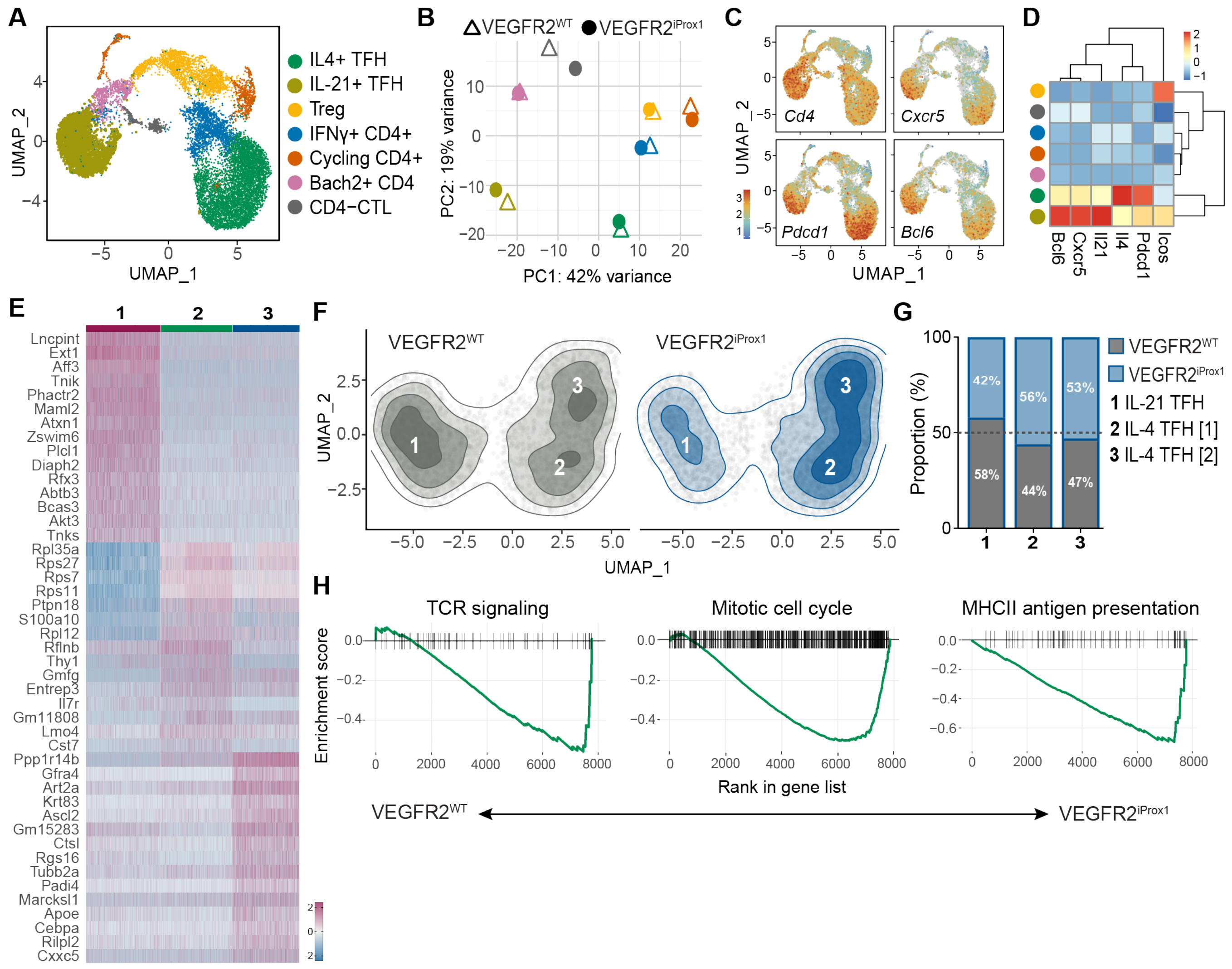


**Supplemental Figure 7. scRNA-seq clustering and transcriptional classification of $T_{FH}$ cell subsets in VEGFR2$^{WT}$ and VEGFR2$^{iProx1}$ mice.**

**(A)** Unsupervised Seurat clustering of UMAP projection of scRNA-seq profiles from sorted follicular helper T cells ($T_{FH}$) (TCRβ$^{+}$B220$^{-}$CD4$^{+}$PD1$^{+}$CXCR5) isolated from dLNs of VEGFR2$^{WT}$ and VEGFR2$^{iProx1}$ mice at day 25 post–VACV infection (by scarification).

**(B)** PCA plot of T cell clusters from (A). Symbol color denotes subcluster, symbol shape denotes genotype. Triangles indicate VEGFR2$^{WT}$ and circles indicate VEGFR2$^{iProx1}$.

**(C)** Expression of key marker genes used to define $T_{FH}$ clusters from total T cells in (A).

**(D)** Heatmap with dendrogram showing $\log_2$ fold changes of hallmark $T_{FH}$ genes and their relatedness between T cell clusters identified in (A)

**(E)** Heatmap comparing gene expression in single cells across 3 identified $T_{FH}$ clusters based on DE analysis included (i) min.pct = 0.2, (ii) min.diff.pct = 0.10, (iii) padj < 0.01. Heatmap describes the top 15 upregulated and 15 downregulated genes for each subset. Color scale indicates $\log_2$ fold change of gene expression. Each column represents a $T_{FH}$ subset, and each row shows z-scored genes expression.

(**F**) Contour plot representing cell density overlay of $T_{FH}$ UMAP, split and colored by genotype (VEGFR2$^{WT}$, grey and VEGFR2$^{iProx1}$, blue).

(**G**) Proportional distribution of each genotype per cluster (VEGFR2$^{WT}$, grey and VEGFR2$^{iProx1}$, blue).

**(H)** Gene set enrichment analysis (GSEA) of TCR signaling, mitotic cell cycle, and MHC class II antigen presentation gene signatures in VEGFR2$^{WT}$ vs VEGFR2$^{iProx1}$.
